# High-throughput DNA repair monitoring in *Saccharomyces cerevisiae* reveals SSB- and DSB-induced chromatin reconfiguration

**DOI:** 10.1101/2024.08.23.609327

**Authors:** Yael Shalev Ezra, Alon Saguy, Gaia Levin, Lucien E. Weiss, Onit Alalouf, Yoav Shechtman

## Abstract

DNA repair is critical for cellular function and genomic stability across organisms. Yeast mating-type switching serves as an established model for studying DNA break repair and chromatin dynamics. However, real-time tracking of mating-type switching in live cells remains challenging due to resolution limitations of existing techniques. Here, we use high-throughput methods, including three-dimensional imaging, to follow the dynamics of DNA damage and repair and to quantify mating-type switching occurrences at the single live cell level, with unprecedented resolution. We reveal chromatin reconfiguration for both single- and double-strand breaks following switching induction. Our findings provide new observation of the correlation between chromatin folding and single-strand breaks.

## Introduction

The spatial organization of nuclear chromatin is nonrandom^1–5^, playing a crucial role in gene transcription, DNA replication, DNA breakage repair^6^, and more. These functions require dynamic conformational changes in DNA^2,3,7–9^. DNA lesions, particularly single-strand breaks (SSBs) and double-strand breaks (DSBs), are genotoxic. SSBs can lead to DSBs in proliferating cells, and DSBs, if left unrepaired, are extremely deleterious events that cause genomic instability^10,11^; thus, DNA damage repair mechanisms are crucial for genome integrity. Homologous recombination (HR) is one of the DSB repair mechanisms. During HR, the DNA lesion is resected to single-strand DNA (ssDNA) overhangs, facilitating proximity between the DSB site and a homologous sequence^12^ which serves as a template for repair. This homologous site is located on either a homologous chromosome, sister chromatid or an ectopic location^11,13–15^. Although repair-related chromatin dynamics have been studied in the past^16–19^, there are still ambiguities regarding the interplay between genome stability, repair and chromatin reconfiguration.

Mating-type switching in *Saccharomyces Cerevisiae* (*SC*) yeast is used as a model system to study HR in eukaryotic cells^11,15^. During mating-type switching, yeast cells switch between ‘a’ (MATa) and ‘α’ (MATα) mating types, to allow sexual reproduction and diploidy^15^. The difference between the two mating types is in the set of genes present in the MAT locus on chromosome III. The set is converted from one type to the other by means of an HO-endonuclease mediated DSB at the MAT locus, followed by repair by HR, i.e. gene conversion, using silent (unexpressed) templates on the opposite ends of the same chromosome, namely HMLα and HMRa^20^. The site-specificity and amenability to genetic modifications has made mating-type switching in *SC* a standard model system for studying DSB repair, shedding light on repair-related chromatin dynamics and biophysical properties in live cells^21–23^. Observation of DNA dynamics during mating-type switching have been previously performed by fluorescently labeling the HMLα and MAT loci with fluorescent repressor-operator systems (FROSs)^24,25^, and measuring their positions over time^21,22,26^.

FROS enables spatially compact fluorescent labelling of the DNA near specific loci, manifesting as a bright spot, and using orthogonal systems enables multichannel observation of several points separately. Experiments using FROSs that suggested contact, repair by gene conversion, and switching, were done either by laser scanning confocal microscopy in fixed cells^21,26^, or by z-scan widefield fluorescence microscopy with very few live cells at low spatial resolution (for example, using a shared fluorescent protein for labeling two loci, which limited spatial co-localization precision)^21,22^. Notably, three-dimensional (3D) positions of fluorescent foci can be determined from the image; the axial position can be found by z-scanning the sample^22,26^, or by point-spread function (PSF) engineering, where an optical element, i.e. a phase mask, is inserted into the microscope, encoding the z position into the shape a point-source generates on the detector^27,28^. This makes it possible to record multiple loci simultaneously and measure their absolute 3D distances^29,30^, at high spatiotemporal resolution. Importantly, in order to obtain robust statistics and overcome biological and experimental noise factors, high-throughput is essential; thus, high-throughput 3D imaging is at the heart of this work.

In this study, we track the 3D DNA dynamics of mating-type switching in live cells at high resolution and high throughput. We use qPCR and flow cytometry to quantify rare switching events in real time and monitor DNA damage repair. Using 3D imaging flow cytometry (3D-IFC), we observed DNA movement during switching induction at unprecedented temporal resolution, and detected locus repositioning within 10 minutes. Our results suggest that upon DNA damage the distance between the fractured and the template loci shortens, regardless of the break type, namely SSB or DSB. This work highlights the power of high-throughput and high-resolution imaging in unraveling the spatial dynamics and reconfiguration of chromatin during DNA damage repair.

## Results

Our objective of tracking DNA dynamics within live cells necessitates precise three-dimensional imaging at high-throughput as well as a robust quantitative mating-type switching assay. To this end, we optimized real-time DNA damage and repair quantification methods based on either qPCR or flow cytometry, each offering distinct advantages. Next, we integrated fluorescent localization systems designed for monitoring chromosomal loci movement and assessed their impact on the mating-type switching process. Finally, we utilized PSF-engineered image-based flow cytometry to observe thousands of cells and measured the 3D spatial distances between the MAT and HMLα loci.

### Quantification of mating-type switching in live cells

To enable live-cell monitoring and quantification of the mating-type switching process, we optimized two methods: qPCR and a fluorescent reporter assay. First, we constructed HO-CFP plasmids designed for both DSB induction and fluorescent reporting. These plasmids contain the HO-endonuclease gene under the GAL10 promoter to induce DSB and CFP under the MFα2 promoter to report switching from MATa to MATα (see methods; figure S1A). For qPCR we built a calibration curve using known mixtures of MATa and MATα cells (see methods and figure S1-B-D). For the fluorescent reporter method we utilized CFP expression, compatible with both FC and IFC. Our HO-CFP plasmids hold several advantages compared to similar plasmids constructed previously^22^: 1. reduced HO-endonuclease background expression - the HO-endonuclease is expressed under a weaker promoter^31,32^ (GAL10 rather than GAL1) which minimizes potential HO leakage, 2. stable expression - the coding genes are stably expressed (integrated into the genome), and 3. enhanced signal-to-noise ratio (SNR) - thanks to the localization of the fluorescent reporter into the nucleus by a nuclear localization sequence (NLS), and due to fusion of 6 msCFP units in some of our strains (see methods).

Throughout this work, we used the BY4741 strain with a “stuck” mutation (*MATa-stk*) at the MATa locus. This strain has a 1 base-pair substitution (T→A at Z11) in the HO-endonuclease recognition site outside of the Ya sequence within the MAT locus, which causes inefficient DSB activity, and accumulation of heterogenous population of cells experiencing either SSB or DSB upon switching induction (figure 1A). This strain is well-suited for tracking DNA dynamics due to the minimal leakage of switching by the GAL10 promoter prior to induction; namely, switching prior to induction is minimal, although its switching efficiency is reduced^15,33,34^.

**Figure 1.**
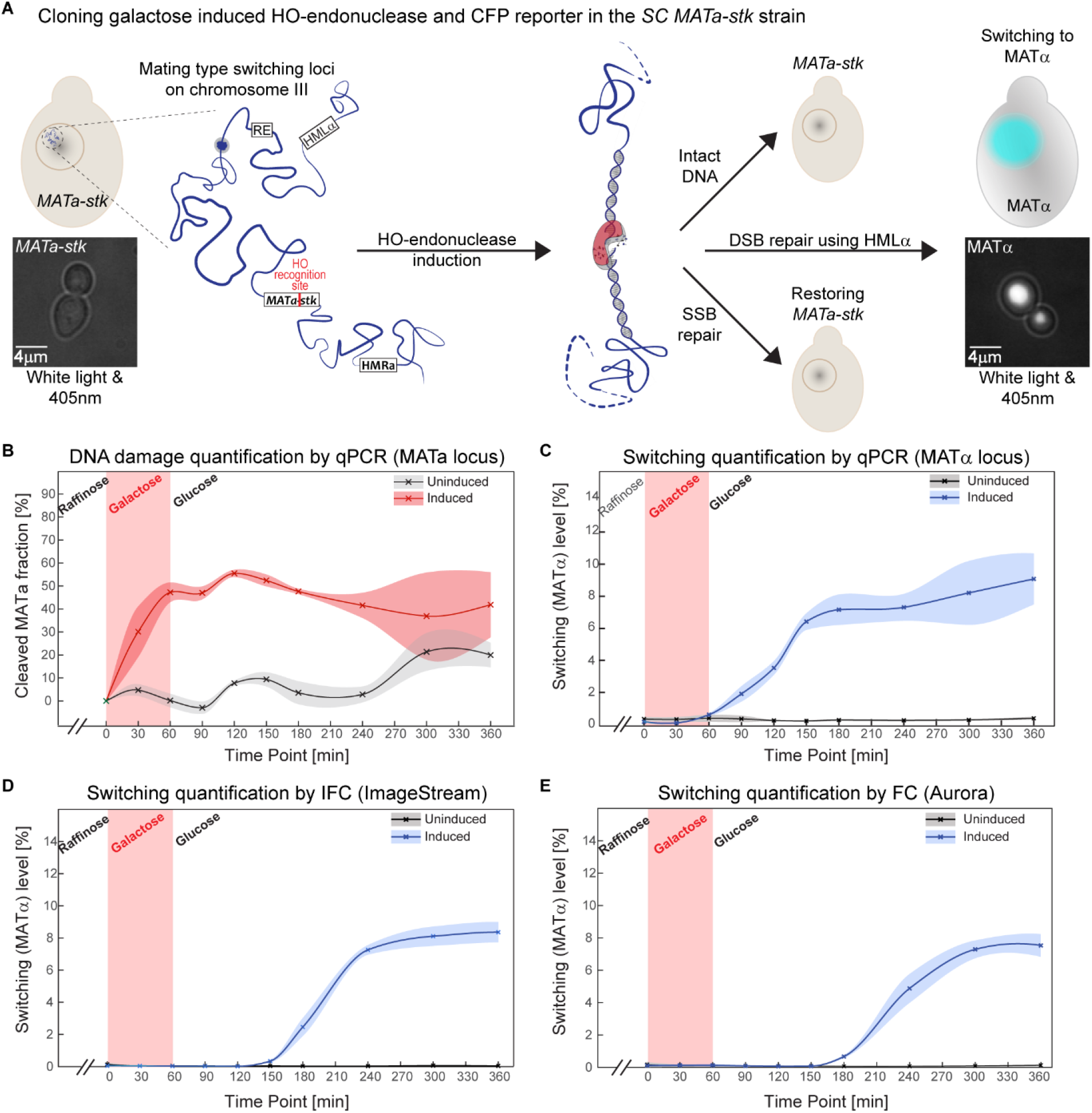
Real-time tracking and quantification of *MATa-stk* switching to MATα. (A) Mating-type switching *MATa-stk* strain, stably expressing inducible HO-endonuclease (by galactose), and the switching fluorescent reporter CFP. Induction of HO-endonuclease results in a mixed population with either intact DNA or damaged DNA (SSB or DSB). Cells that repaired their DNA by switching, express CFP in the nucleus. (B) *MATa-stk* locus quantification by qPCR; disappearance of this locus quantifies the DNA damage, while its reappearance corresponds to non-switching repair. Tracking and quantifying switching by (C) qPCR of MATα locus, and CFP as fluorescent marker in (D) IFC, and (E) FC. The ~90 minute difference between switching detection by qPCR (C) vs. flow methods (D),(E), is consistent with CFP maturation time of ~90 minutes (see table S1). All experiments obtained significant switching levels (*p-value<<0*.*01*) by comparing induced to uninduced (0% galactose) cultures using either N-way or two-way ANOVA.

We transformed the *MATa-stk* strain with the switching and reporting plasmid (figure S1A) and induced mating-type switching in liquid culture, after initial growth in raffinose, by the addition of galactose (2-4% w/v) to the media, prompting HO-endonuclease expression. To terminate HO-endonuclease expression, which has a short half-life time of ~10 minutes^15^, we added glucose (2% w/v) after 1 hour of induction (see methods). To validate the HO-CFP applicability, we quantified switching after 6 hours from induction both by qPCR and by imaging the nuclear accumulation of the CFP reporter (figure 1A), and showed consistency between the two methods among different colonies with different switching levels (figure S1E)^33,34^. Additionally, we found our HO-CFP plasmids to induce switching more efficiently compared to pJH132^35^ and B2609^22^ centromeric plasmids (figure S1F). See supplementary section S1 and supplementary figure S2 for validation of the switching efficiencies.

Next, we monitored the mating-type switching process for 6 hours, collecting samples every 30-60 minutes to quantify DNA damage and switching levels. Using qPCR with primers that span both part of the unique Ya region and the HO-endonuclease cut site in the MAT locus (figure S1-B) we followed DNA damage occurrence (figure 1B), including repair back to *MATa-stk* type, i.e. repair that is not associated with switching (figure S3-A); additionally, we quantified the switching level in the population using both qPCR of the MATα (figure S1-B) and CFP detection. The qPCR measurement of MATa detected ~30% DNA damage after 30 minutes from induction, which increased to over 50% after 120 minutes (figure 1B). The decrease in DNA damage of MATa to ~40% starting from 120 minutes after induction till the end of the experiments (figure 1B), indicates that at least 10% of the population, with any type of DNA damage, did not switch to MATα, and repaired the damaged DNA back to *MATa-stk* (figure 1B and figure S3-A).

To quantify switching more directly, we performed qPCR with MATα primers and measured CFP expression in live flowing cells. The result shows that switching occurred already as soon as 60 minutes after induction (figure 1C-E), earlier than implied previously (70 and 105 minutes^36,37^ as measured by qPCR and southern blot). Different concentrations of galactose for induction (2% and 4%; data is shown for 2%) resulted in consistent behavior across these independent measures, namely, qPCR (figure 1C) and CFP monitoring by IFC (figure 1D) and FC (figure 1E), overall revealing a 6-10% switching rate at the end of the experiments. The rest ~30% of the population, which were neither the ~60% *MATa-stk* type nor the 6-10% switched cells (MATα), probably did not repair the DNA damage at all, in agreement with previous studies^13,38,39^. Overall, these results imply that the switching limiting step in the strains we used (BY4741 derivatives) is due to the “stuck” mutation in the MATa locus that reduces the DSB probability in the population, which agrees with previous studies^15,33^.

Interestingly, using flow cytometry, where switching was quantified by CFP level, we observed a negative correlation between the switching ratio and the relative number of singlets (i.e. single cells; figure S3-B); This can be explained by the fact that as the population underwent switching, mating events followed, manifested as doublet (non-singlet) events, which were observed in the forward and side scattering-channels. This suggests that, under some conditions, switching levels may be quantified without need for a fluorescent reporter at all. Indeed, calculating the fraction of switched cells expressing CFP in the singlet and in the non-singlet populations showed that non-singlet cells expressed CFP with significantly higher probability than singlets (figure S3-C-D). We can therefore approximate the switching level (*Sw*) by:

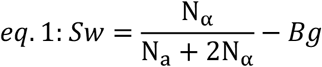

where N_*α*_ is the number of MATα cells, N_a_ is the number of *MATa-stk* cells, and *Bg* is the non-singlet background level calculated using the uninduced culture. The calculation above assumes that each switched cell (MATα) partners with a single unswitched cell (*MATa-stk*), manifesting as a non-singlet reading in the flow cytometer, which we find to be a good approximation in our case, supported by the fact that the calculated switching levels over time (figure S3-E) were similar to the qPCR, FC and IFC measurements. For example, at 300 minutes post induction, using this label-free method we calculated 6.72% switching in the population, whereas 8.21%, 7.29% and 8.11% were estimated using qPCR, FC and IFC, respectively. Thus, simply counting singlets vs. non-singlets with a flow cytometer is useful as a label-free method to estimate mating-type switching via mating in *MATa-stk* cells. This detection method has a limited time frame: as the culture grows, aggregates form naturally, independent of mating events. This aggregation was visible in the uninduced culture plot, where there was a slight reduction in the number of singlets (figure S3-B), despite the lack of switching.

To summarize this part, we have optimized several quantitative methods for mating-type switching detection. The choice of method ultimately depends on the experimental requirements. For example, the CFP-based methods are live-cell compatible, sensitive, and successfully report on the rare switching events during the experiment, on the single-cell level, however detection is lagging by approximately 90 minutes due to the fluorescent protein expression and maturation time^40^. The qPCR method, on the other hand, detects switching after 60 minutes, however, it requires a tedious sample preparation procedure, including a genomic DNA extraction step. The different attributes of the methods are summarized in table S1.

### Integrated fluorescent repressor-operator system (FROS) for live-cells DNA dynamics

For live cell imaging we used FROS^8^ to generate observable fluorescent spots near the HMLα and the *MATa-stk* loci (figure 2A) in a mating-type switchable strain, and measured the distances between the two localizations over time. The loci are expected to be in close proximity after DSB induction for the HMLα to serve as a gene conversion template^15,22,26^. FROS is a highly useful tagging approach for specific loci tracking in live cells, despite its limitations related to fluorescent background and spontaneous loss of operators^41^. Specifically, when applying PSF engineering microscopy, such as in our 3D IFC measurements (see next section), high SNR is important because photons are spread over a larger area compared to the standard PSF^29,30^. Thus we optimized new strains, rather than using existing ones^23,26^, for fluorescent loci with high SNR. We found that eGFP (near the *MATa-stk* locus; LacO, LacI-eGFP) and mCherry (near the HMLα; TetO, TetR-3xmCherry) outperformed YFP and mRFP^23^ with regard to SNR and spectral differentiation. Furthermore, to improve localization precision we screened for colonies with the highest SNR (see methods and figure S4).

**Figure 2.**
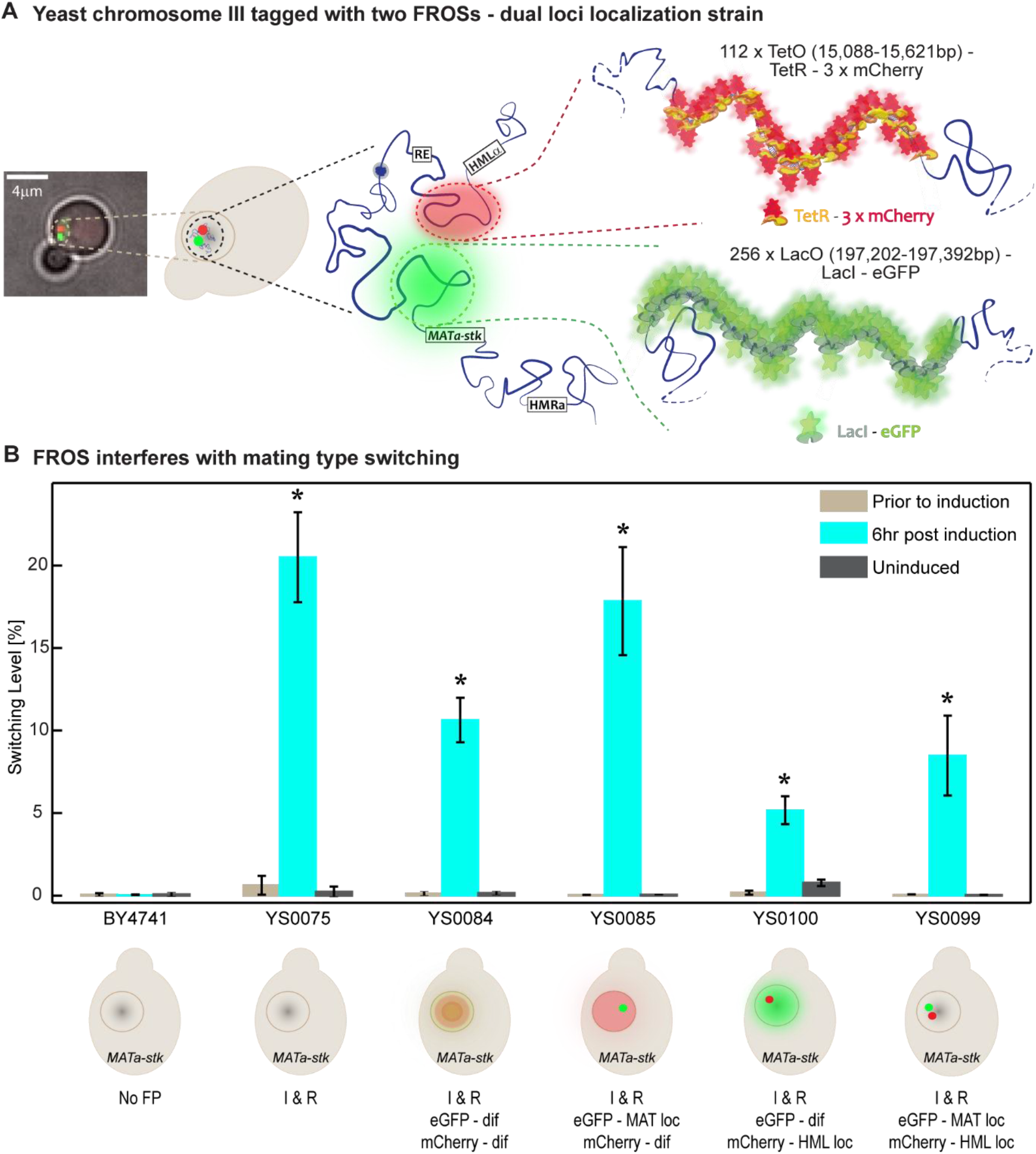
Integrated FROS reduces switching efficiency. (A) Example image and schematic illustration of two FROSs labeling the mating-type switching loci in live yeast cells. Homology regions for integration are specified in parentheses. (B) The effect of FROS genome editing on mating type switching efficiency. The frequency of switching is significantly influenced both by genomic perturbations next to the mating-type switching related loci, and by the diffusing fluorescence repressors (*p-value<<0*.*01*; t-test). Abbreviations: I & R - Inducible and reporting strains containing the HO-CFP plasmid; dif - diffuses in the nucleus; loc – localized in specific locus (HMLα or MAT).

Notably, FROS integration into the yeast genome is expected to cause chromatin disruption, such as folding abnormalities^41^; therefore, since chromatin reconfiguration is key in mating-type switching, we quantified the effect of FROS on switching efficiency. We compared switching efficiencies between HO-CFP strains that contained diffusing fluorescent proteins (i.e. no operator repeats), a single FROS, and two FROSs. We found that insertion of the FROS indeed adversely affected the repair process during gene conversion and significantly reduced switching efficiencies; however, interestingly, while the operators of the TetO-TetR-3*mCherry (next to HMLα) caused reduced switching occurrence, in the LacO-LacI-eGFP system (next to MAT) the reduced efficiency occurred mainly due to the free LacI-eGFP molecules, and the LacO-LacI-eGFP operators had a very minor effect (figure 2B). Overall, switching efficiencies were reduced by 2-3-fold, due to both the diffusing fluorescent proteins and the operator repeats.

### Population analysis of DNA repair dynamics using 3D IFC reveals SSB- and DSB-related DNA reconfiguration

Combining flow cytometry with three-dimensional (3D) nanoscale resolution and multi-color fluorescence imaging enables high-throughput data collection, and sub-nuclear visualization with high precision and accuracy in single live cells, on the population level. We used a modified 3D IFC system^42^ (figure 3A-D); a cylindrical lens inserted into the IFC generates an astigmatic PSF that encodes the three-dimensional positions of the loci, enabling 3D distance measurement from a snapshot, at 60-80 nm colocalization precision (see methods and figure S5-A).

**Figure 3.**
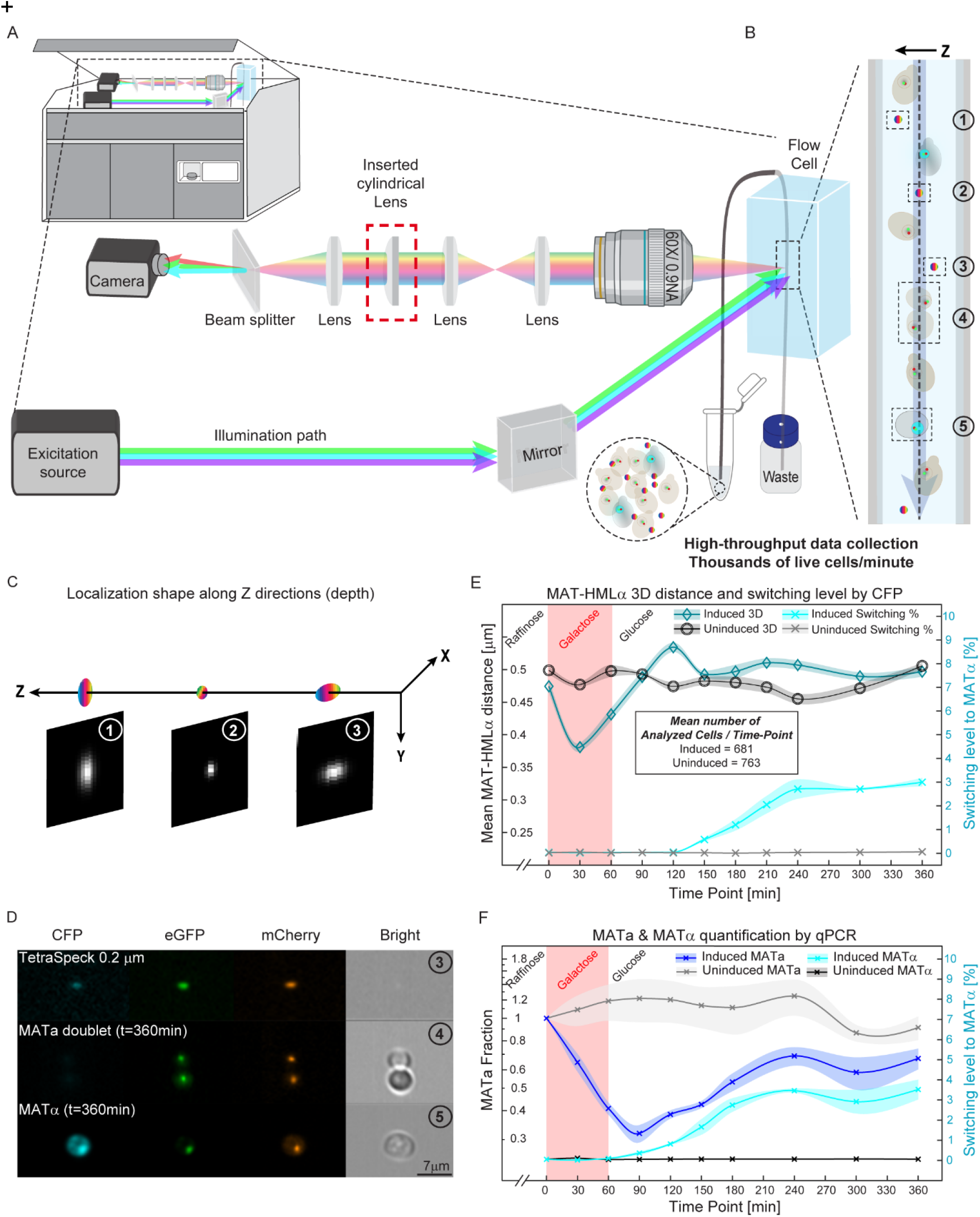
High throughput tracking of *MATa-stk* and HMLα loci dynamics (mean distance) during switching. (A-D) IFC instrument setup and imaging: (A) a cylindrical lens is incorporated into an ImageStream IFC, for 3D acquisition; (B) beads were added to each of the culture samples for simultaneous 3D position calibration at each time point of the switching experiment; (C) example images of bead PSFs at different 3D positions in flow (beads 1, 2, 3 in (B)); (D) image acquisition of a bead and cells in 4 channels: detection of switching occurrence in the CFP channel, 3D localizations of MAT and HMLα foci in the eGFP and mCherry channels, respectively, and object appearance in brightfield channel. (E) Real-time 3D distance measurements between MAT and HMLα and switching levels by CFP detection of an induced and uninduced cultures. In E 3 replicates were analyzed via bootstrapping: mean (line) ± s.d. (shaded region representing the estimated error) of eight independent groups in each time point, overall >429 cells were analyzed per time point. Distance shortening and switching were detected at 30 and 150 minutes, respectively, post DSB induction (*p-value* << 0.05). (F) DNA damage and switching levels for the dually tagged strain (YS0099) by qPCR (*p-value* < 0.01; 3 biological replicates). Statistical analyses were performed with two-way ANOVA.

The high throughput of IFC is crucial for the ability to obtain statistically significant results. We were able to image thousands of cells per minute and analyze only cells with good SNR spots (5-10% of the imaged cells), maintaining statistical significance despite low switching levels. Moreover, employing 3D rather than 2D analysis provides greater precision for assessing DNA loci proximity; which is especially relevant for short distances such as in the yeast nucleus that has a diameter of approximately 1.5 µm (depending on the carbon source^42–44^).

Initially we measured the native 3D distances between the HMLα and *MATa-stk* loci, which ranged from 300 to 500 nm across the same strain obtained by different transformations (figure S5-B). This variability may be attributed to the inherent heterogeneity in chromatin configuration^7,41,45^ and is further influenced by the perturbation obtained through FROS integration (FROS integrations were verified by sequencing). We then proceeded to the switching experiments, acquiring images every 30 minutes. We observed significant change in proximity (mean distance) between the fractured *MATa-stk* locus and the HMLα template both in 2D (figure S5-C) and 3D (figure 3E, figure S5-D) as soon as 30 minutes after induction, earlier than detected before (80 and 60 minutes^22,26^). Similar analysis we performed by continuous monitoring revealed that the change in proximity due to induction of DNA damage and switching starts as early as 10 minutes after the induction (see supplementary section S3 and figure S6). Despite the fact that the native distance between MAT and HMLα was larger in MATα than in MATa (figure S5-E and in accordance with previous studies^23,46^), we did not observe significantly larger mean distance in the switched population at the end of the experiment. This is to be expected, due to the low switching occurrence (figure 3E).

According to our qPCR results on the dually-tagged strain, while 68% of the population experienced a DNA damage event, (within 90 minutes), at the end of the experiment only ~3.5% switched to MATα, thus underwent DSB and used HMLα as a template for repair (figure 3F, consistent with the CFP measurements in figure 3E). Additionally, 36% repaired back to *MATa-stk* type. The rest of the population (~30%) that was detected neither as MATa nor MATα, probably did not repair the damaged DNA at all (figure 3F; similar to figure S3-A).

In accordance to our observation in the previous section, FROS interfered with the mating-type switching process; this can be seen from the fact that although the dually FROS-tagged strain experienced ~15% more DNA damage compared to strain with no FROS (compare figure 3F to figure S3-A), we observed ~2.5-fold less switching to MATα (3-3.5% in figure 3F vs. 6-10% in figure 1C), and approximately 3.6-fold more cases of repair back to *MATa-stk* (36% in figure 3F vs. 10% in figure S3-A).

The discrepancy between the low switching levels (3-3.5%) and the striking distance shortening of ~120 nm (from 470 to 350 nm) between HMLα and *MATa-stk*, observed 30 minutes after induction on the population level (figure 3E), calls for an explanation. Clearly, attributing this distance shortening only to switched cells is not sufficient; this can be shown by calculating the expected mean inter-loci distance in the population, even under the extreme assumption that 3.5% of the population is at distance 0 (i.e. the two loci are touching), at time points 30 and 60 minutes. The result is shown in figure 4A, on the uninduced population, exhibiting, as expected, a negligible expected effect on mean distances.

**Figure 4.**
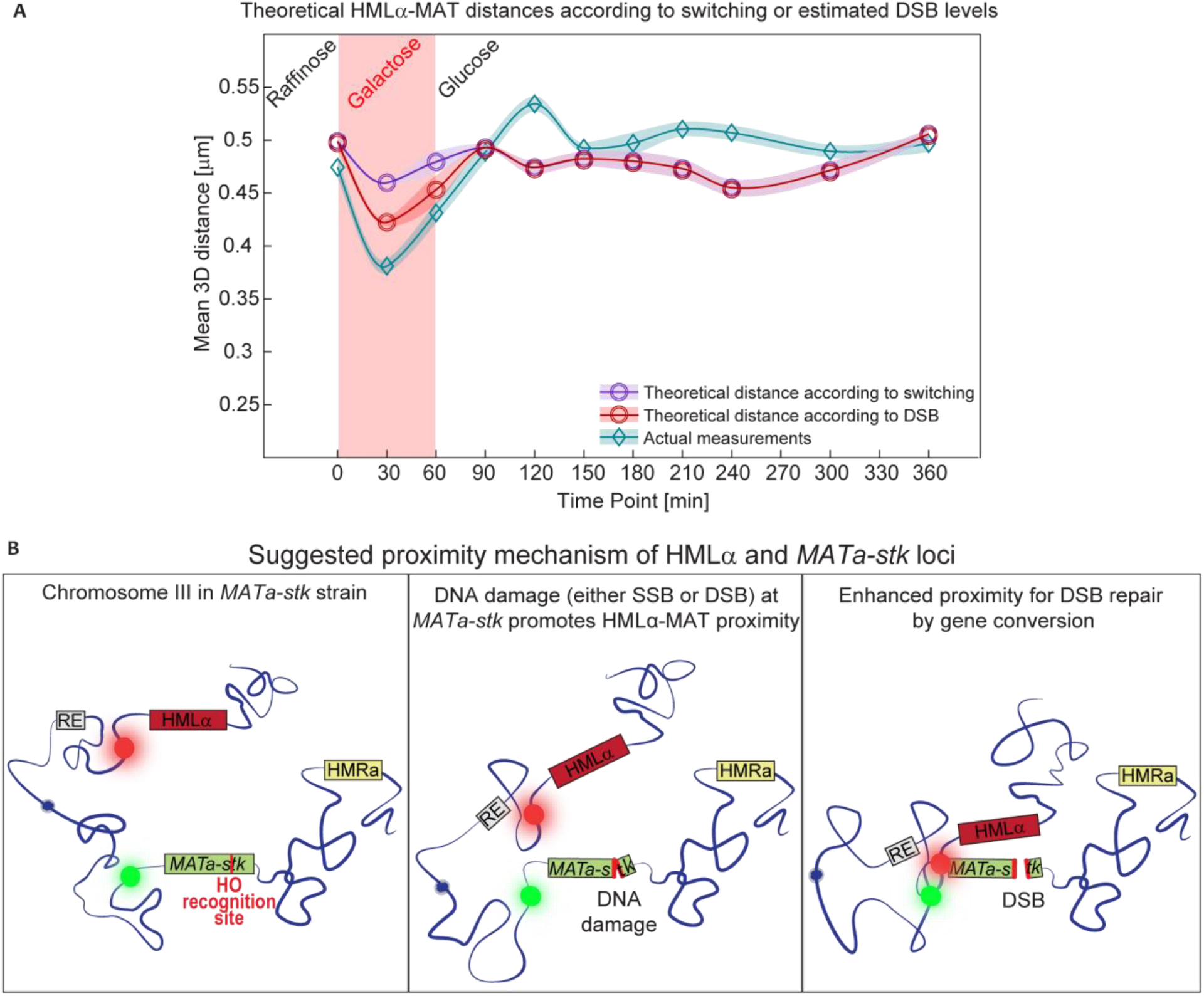
DNA damage at the MATa-stk locus results in proximity between the HMLα and MAT. (A) Hypothetical 3D HMLα-MAT distances using uninduced 3D IFC data; distance zeroing of 3.5% (switching) or 10% (DSB) of the population at 30- and 60-minutes post induction. (B) Suggested model for HMLα and *MATa-stk* dynamics upon switching induction: intact chromosome III configuration (left); *MATa-stk* locus cleavage by HO-endonuclease results in directed movement bringing HMLα and *MATa-stk* closer together (middle); Further movement and closer proximity occurs in cells with DSB (but not SSB) at the *MATa-stk*, which promotes contact between HMLα and MAT, repair by gene conversion and switching (right).

Since switching rate is not high enough to explain the substantial level of HMLα-MAT proximity shown in the induced strain (figure 3E), we hypothesize that this effect is caused by DNA damage, whether resulting in switching or not. One possibility would be that the observed proximity in inter-loci distance is associated with DSB. To explore this option, we repeated our analysis, namely, zeroing the inter-loci distances of a subset of the population that corresponds to the fraction of cells exhibiting DSB, and checking whether the proximity-dip shown in figure 3E is explainable this way. Since we do not know the DSB rate directly, we estimate it using a strain without FROS (based on the results in figure 2), where switching was 2-3-fold higher than the strain with FROS, and assuming that each DSB corresponds to a switching event in the strain without FROS; even when considering this rate of about 10% DSB (calculated as ~2.8 *×* 3.5%) at time points 30 and 60 minutes, there was still an unexplained proximity between the loci in the population (figure 4A). Since most of the population experienced DNA damage, either SSB or DSB according to our qPCR results, we finally conclude that the HMLα-MAT proximity is caused both by DSB and by SSB.

This analysis suggests the following dynamic model (figure 4B): after an induced DNA damage (either DSB or SSB) in *MATa-stk* by the HO-endonuclease, there is DNA motion which promotes proximity between the *MATa-stk* and HMLα loci, regardless of the type of damage. DSB events will promote further directed DNA movement and physical association that allow for lesion-template hybridization and DSB repair by gene conversion, namely, mating-type switching.

## Discussion

In this work we presented high throughput methods for mating-type switching investigation in live yeast cells bearing the *MATa-stk* mutation. We used advanced biological, optical, and analytical tools to detect and follow switching occurrence, DNA damage, and related DNA dynamics with high precision and single cell sensitivity, and revealed new findings, detailed below.

Firstly, using quantitative real-time single live-cell analysis of switching occurrence with high throughput, we were able to detect rare switching events in a heterogenous population. Next, we were able to quantify and unveil the influence of FROS tagging on the mating-type switching process, which interrupts DSB repair. Moreover, we showed that FROS, which serves as a popular DNA tagging for *in-vivo* tracking^47,48^, is still a valuable tool here, as its detrimental effect on switching rate can be compensated by using high throughput methods like those presented here. Additionally, we found that different transformants of the same strain exhibited different spatial conformations of chromosome III, as indicated by differences in the native inter-loci distances we measured.

Secondly, we demonstrated the potential of the high throughput 3D-IFC to study DNA damage repair dynamics. We monitored DNA dynamics with 2-3 fold higher spatial precision than previously reported (60-80 nm compared to 200-300 nm^21–23,46^). Importantly, this high throughput approach allows imaging of thousands of live cells, selected for optimal SNR, to observe mating-type related loci during switching with high confidence, compared to fixed cells by confocal microscopy^26^. Taken together, we report two new main findings upon mating-type switching induction: 1. A motion that brings HMLα and *MATa-stk* loci closer to each other as soon as 10 minutes after induction, which suggests to be independent of the gene conversion repair process, which starts after more than 10 minutes^11,49^; and 2. The motion occurs in all cells with DNA damage at the *MATa-stk* locus (either SSB or DSB), however, tighter proximity between the loci seems to correlate only with DSB.

The observable proximity between the HMLα and MAT during DNA damage at the *MATa-stk* locus suggests the underlying mechanism: upon DNA damage (SSB or DSB) the two loci approach each other, possibly facilitated by the cis-acting element, namely the recombination enhancer (RE), and related proteins, such as the Fkh1 protein^15^. The Fkh1 protein has multiple binding sites on the RE and contains a conserved amino acid sequence, known as the FHA domain, that interacts with phosphorylated DSB repair proteins during the mating-type switching procecss^15,50–52^. Based on our finding that switching and DSB rates are too low to explain the observed inter-loci proximities, and on previous works in mammalian cells^10,53^ and yeast cells^50,51,54^, it is possible that Fkh1 binds phosphorylated SSB repair proteins near the damaged site as well^10,53–55^. This suggests that the RE may respond not only to DSB but also to SSB repair mechanisms.

In cells with DSB (but not in cells with SSB), a closer proximity increases the probability of switching and likely involves proteins which are relevant only for DSB repair^11,15^. This tight proximity may include multiple and reversible contact events between the HMLα and MAT loci in each induced cell^22^, that are interrupted in the dually tagged strain with the FROSs (figure 2B). Further experiments with gene conversion related tagged proteins, such as CK2, Fkh1 or Rad51, and their effects on DNA dynamics during the switching process will enhance the understanding regarding the contribution of each protein in this process.

## Methods

### Plasmid construction

Plasmids used in this study are listed in table S2. The new plasmids were constructed by Gibson assembly^56^, or by standard restriction and ligation using T4 ligase (M0202, NEB). Gibson fragments were amplified by polymerase chain reactions (PCRs) using either the PrimeSTAR GLX (R050A, Takara), LA Taq Hot Start (RR042A, Takara) or Q5 DNA (M0491, NEB) polymerase, with the appropriate primers (listed in table S3). All homologous recombination sites were amplified from *S. cerevisiae* (BY4741) genomic DNA, their locations are specified below. All plasmid modifications were validated by sequencing.

pYS0080 (CyPet expression under MFα2 promoter) was assembled by multiple Gibson steps using pUC19 backbone and DNA fragments amplified by primers P-YS1 to P-YS12 (pCyPet-His was a gift from Patrick Daugherty (Addgene plasmid # 14030^57^); pYTK029 was a gift from John Dueber (Addgene plasmid # 65136^58^); BS-Met15 was a gift from Zhiping Xie (Addgene plasmid # 69198^59^)); and the HR site was amplified from ChrXII (730361-730807 bp) with P-YS11 and P-YS12 primers. pYS0088 was assembled with pYS0080 backbone and CFP that was amplified from pIL01^23^ by PCR using P-YS13 and P-YS14 primers. pYS0099 was assembled with pYS0088 backbone and HO-endonuclease under the control of GAL10 promoter, namely, GAL10p-HO, which was amplified from pJH132^35^ by PCR using P-YS15 and P-YS16 primers. pYS0107 was constructed from pYS0099 backbone, which was amplified by PCR with P-YS17 and P-YS18 primers, and 6XmsCFP, which was extracted from YIplac211-SEC7-msCFPx6 (a gift from Benjamin Glick (Addgene plasmid # 105268)^60^), using EagI-HF (R3505, NEB) and NheI-HF (R3131, NEB) restriction enzymes.

pYS0019, carrying 112 TetO repeats with URA3 as a selection marker, was constructed from dually digested pKW2837^8^ with SacI (R3156, NEB) and NgoMIV (R0564, NEB) and HR site proximal to HMLα (15,088-15,621 bp, using P-YS19 and P-YS20 primers). pYS0020, carrying 256 LacO repeats with LEU2 as a selection marker, was constructed from dually digested pKW1689^8^ with Acc65I (R0599, NEB) and XhoI (R0146, NEB) and HR site proximal to HMRa (294,898-295,239 bp, using P-YS21 and P-YS22 primers). pYS0021 was constructed from dually digested pYS0020 with Acc65I and XhoI by replacing the HMRa HR site with HR site proximal to MAT (197,202-197,392 bp using P-YS23 and P-YS24 primers).

### Yeast strain construction

All strains (listed in table S4) were derived from BY4741 and BY4742^61^, non-homothallic strains carrying mutated *ho*. BY4741 strain possesses the *MATa-stk* mutation, a single-base-pair substitution in the HO-endonuclease recognition site (T to A in Z11^15,33^). All strains were constructed by transforming linear plasmid (listed in table S2), digested by the appropriate enzymes, using the LiAc/SS carrier DNA/PEG method^62^. The transformed strains were grown on the appropriate selectable medium, validated by colony PCR (with primers listed in table S3), sequencing and by microscopy (figure S3-A). We optimized our experiments by screening for colonies with the highest signal-to-noise ratio (SNR), using the IFC; we compared the SNR of each localization to a strain expressing the fluorescent proteins with no operator repeats (low SNR), and to 0.2 µm TetraSpeck beads (TS200; T7280, Invitrogen) (high SNR). The colonies with the higher fraction of cells above the SNR threshold (equals to the mean of colony SNR means), and with two visual bright loci observed by IFC and fluorescence microscopy, were chosen (figure S3-B).

The inducible and switching reporting strains YS0075 and YS0095, were constructed by transforming BY4741 with linear pYS0099 and pYS0107, respectively.

For dual-color tagging of the HMLα and MAT loci, we used the LacI-LacO^24^ and TetR-TetO^25^ FROSs, respectively. First, YS0007, YS0020 and YS0084 were constructed by transforming BY4741, BY4742 and YS0075, respectively, with NheI-HF linearized pKW3034 plasmid (TetR fused to 3XmCherry and LacI fused to eGFP)^8^. YS0104, YS0057 and YS0085 were constructed by transforming YS0011, YS0051 and YS0084, respectively, with PmlI linearized pYS0021 plasmid (256xLacO repeats). YS0011, YS0051, YS0099 and YS0100 were constructed by transforming YS0007, YS0020, YS0085 and YS0084, respectively, with BmgBI linearized pYS0019 plasmid (112xTetO repeats).

YS0096 strain was derived from YS0095 by switching induction and sorting twice CFP expressing cells (first sorting with Bigfoot Spectral Cell Sorter by Invitrogen, and second sorting with the FACSAria™ III Cell Sorter by BD). YS0096 MAT type was validated by colony PCR (with two sets of primers for the amplification of the MAT locus: P-YS41, P-YS42, P-YS43; see table S3), microscopy, and mating assay.

### Media and growth condition

All experiments started from a single isolated colony grown at 30°C and 200 rpm. For nonselective growth, we used standard YEPD medium (1% yeast extract (w/v), 2% bacto-peptone (w/v) and 2% dextrose (w/v)). For selective growth, we used defined SD medium (0.67% YNB without amino-acids (AA) (w/v) and 2% dextrose (w/v) supplemented with all necessary AA and nucleobases according to strain requirements; as indicated in Cold Spring Harb Protocol^63^. We added 1.5% bacto-agar (w/v) for petri dish growth). For microscopy visualization, cells were grown in liquid defined SC \ SD (−Leu or −Ura) low fluorescent medium (LFM) (0.69% YNB (without AA, folic acid, and riboflavin) (w/v), all necessary AA and nucleobases according to requirements, and supplemented with 2% carbon source (raffinose/ galactose/ dextrose) (w/v)).

Importantly, we used the same medium for the starter and culture to avoid hindrance of the fluorescent protein expression, since we found FROS to be sensitive to media change; when we used YEPD starters for raffinose culture, localizations were poor and, in some cases, invisible.

### Induction of Mating-type switching by galactose

Single isolated colonies were grown in 2 ml SC/SD-2% Raffinose (LFM) media for 2-7 days at 30°C and 200 rpm for sufficient cell density. Cultures were resuspended in fresh SC/SD-2% Raffinose (LFM) in baffled Erlenmeyer flask for additional overnight incubation, to reach optical density at 600 nm (O.D_600_) ~0.25-0.35 (about 10^7^ cells/ml), which was just before entering logarithmic phase at time of induction (t=0); O.D_600_ was measured using Ultrospec10 spectrophotometer (Biochrom). The culture was divided into two baffled Erlenmeyer flasks, for switching induction by galactose (induced culture), and for control without induction (uninduced culture), which were then incubated at 30°C and 200-250 rpm throughout the experiment. Switching was induced by adding stock solution of 20% galactose to the culture (2% (w/v) final concentration in culture), and terminated after 60 minutes (unless specified differently) by adding 40% glucose stock solution (2% (w/v) final concentration in culture). Samples were collected for analysis by IFC, FC, and qPCR before, during, and after induction, at 30-60 minutes intervals and up to 6 hours post induction.

### Mating-type switching analysis by flow cytometry (FC)

Yeast culture samples (200-600 µl) during the switching experiments were analyzed using the Cytek Aurora full spectral FC device. Data analysis was done using the SpectroFlo software. ‘Cells for analysis’ were gated using SSC-H X FSC-H, and ‘singlets’ were gated using SSC-H X SSC-A (figure S7-A-B). CFP expressing cells (switched from *MATa-stk* to MATα) were gated based on the unstained culture (BY4741), the culture before induction (YS0095) and a CFP positive strain (YS0096) (figure S6-C-E).

### Mating-type switching analysis by qPCR

5 ml culture samples were obtained during the switching experiments for genomic DNA (G-DNA) extraction by precipitation as follows: Cells were centrifuged for 5 minutes at 1,377 rcf, harvested and resuspended in the minimal medium left after removal of most of the supernatant; G-DNA was precipitated using MasterPure Yeast DNA Purification Kit (MPY80200) according to the kit protocol with the following modifications: centrifugation steps were done in 4°C, microtubes were left open for 15-20 minutes to ensure ethanol evaporation, and DNA pellets were resuspended in 50-100 µl MBW.

For switching and DNA damage quantification, each G-DNA sample was analyzed by qPCR with three sets of primers (figure S1-B): 1. P-YS41 and P-YS42^23^ for MATα genomic region, 2. P-YS42 and P-YS43^23^ for MATa genomic region, and 3. P-YS44 and P-YS45 for ACT1 region (housekeeping gene). Based on primer calibration assay (figure S1-C), G-DNA samples were diluted to 5 ng/µl. Each qPCR reaction contained 6.8 µl MBW, 10 µl PerfeCTa SYBR Green FastMix Low ROX (2X), 0.6 µl (10 µM) of each primer (forward and reverse), and 2 µl G-DNA (10 ng). We used a standard qPCR cycling protocol (QuantStudio1, Applied Biosystems) with initial denaturation stage (3 minutes, 95^°^C), 40 amplification cycles (15 seconds, 95^°^C), end of extension stage (1 minute, 60^°^C) and a dissociation stage. Raw cycle threshold (Ct) data was analyzed to assess fold enhancement (switching) and reduction (DNA damage) using pfaffl method^64^. To quantify switching levels we plotted a standard curve based on multiple known mixes of BY4741 (*MATa-stk*) and BY4742 (MATα) cultures: stationary phase cultures at the same O.D_600_ were mixed, and immediately their G-DNA extracted to avoid mating events; the samples were analyzed by qPCR with MATα primers, and; the Ct values were used to a generate standard curve in the range of 0.5%-30% MATα (figure S1-D). MATa switching events to MATα were accurately quantified by fitting the Ct values to the standard curve equation. DNA damage levels were assessed with MATa primers (P-YS42 and P-YS43), based on the knowledge that different DNA lesions intrude the polymerase and reduce amplification^65^, which was calculated relatively to the housekeeping gene (ACT1) and the starting point (prior to induction).

### Imaging Flow cytometry (IFC)

#### Instrument setup

High throughput live cell 3D imaging was done using the ImageStream®X Mark II (Amnis) IFC, modified by the insertion of a cylindrical lens (*f*=1m, LJ1516RM-A, Thorlabs), which was mounted by kinematic magnetic bases (KB2×2, Thorlabs) into the optical path as described previously^42^. We used the following device settings: (1) 60X/0.9NA objective, (2) turning off autofocus and determining manually the ideal focal position for each run, (3) operating 3 lasers: 405 nm (120 mW) for CFP detection in channel 1, 488 nm (200 mW) for eGFP detection in channel 2, 561 nm (200 mW) for mCherry detection in channel 4, and (4) operating the brightfield illumination in channel 6.

#### Data acquisition

Induced followed by uninduced live cell culture aliquots (200 µl) were mixed with 0.2 µm TetraSpeck beads (0.2 µl; diluted by 1000), flowed and imaged prior to induction (defined as t=0), every 30 minutes for 4 hours, and after 5 and 6 hours post induction (11 time points in total).

As a first pre-processing step, the brightfield channel helped us to distinguish between bead images and cell images. We then gated images that contained intact singlet and doublet cells (area <80, aspect ratio >0.35), in focus (gradient RMS >50), and high SNR (discarding noisy cells by ‘raw max pixel’ for channel 2 and channel 4). These cells were analyzed for CFP expression by ‘intensity’ and ‘raw max pixel’ in channel 1 for real-time switching occurrence (figure S8-A). Next, we exported the bead images and the high SNR cell images into rif, cif, daf and txt files, beads data for calibration, and cells data, generated by the ISX and the IDEAS software, for 3D distance analysis. The rest of the analysis was done by an automatic analysis pipeline publicly available online, and divided into four main parts: 1) pre-processing of the cif data; 2) calibration of PSF shapes to z positions by beads; 3) 3D localization estimation by Gaussian fitting; 4) 3D distance prediction.

#### Pre-processing and parameter initialization

The first step in our analysis pipeline involved converting the cif file to a tif format. Additionally, we performed mean intensity padding to the maximal image size in the cif data and normalized the pixel intensities per image to be in the range [0, 255]. We repeated this step twice: for the green channel (channel 2), and for the red channel (channel 4).

Next, we initialized some parameters according to our ImageStream device, such as: the pixel size, the diameter of the flow tube, etc. We also determined the minimal signal-to-noise ratio (SNR) that was required from an image to enter the analysis pipeline. We used this threshold since locus localization for images with low SNR was less accurate and hindered the ability of our pipeline to detect small inter-loci distance changes over the cell population.

#### Calibration by beads

To obtain the 3D positions of the loci inside the cells we used a cylindrical lens for PSF engineering. When imaging with a cylindrical lens, the PSFs are gradually stretched as a function of their z position in along different axis above and below the focal plane. To relate each PSF shape (width and stretch direction) to a specific z position, we performed calibration between the PSF shape and the z position of the locus.

We found out that performing calibration by the images of labeled loci inside cells presented challenges due to background noise and low SNR. Therefore, we performed the calibration using fluorescent beads, characterized by their high SNR and lack of cell background noise (figure S8-B).

Each bead PSF was fitted to a 2D asymmetric gaussian model denoted by eq. 2:

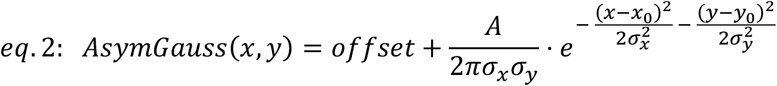

Where: *offset* is the mean background value, *A* is the amplitude, *σ*_*X*_, *σ*_*Y*_ are the gaussian standard deviation estimates, and *x*_0_, *y*_0_ are the bead location estimates.

We drew a histogram of the predicted *σ*_*X*_, *σ*_*Y*_; then, we calibrated the histogram of each sigma to the z position of the bead, namely, we ended up with two calibration functions, one for each axis. Similar to previous work^42^, we assumed the bead position in each axis inside the flow cytometer was approximately normally distributed with 95% of the beads within a cylinder with radius of 2.2 µm. Hence, we could fit each sigma histogram to normally distributed positions in the z axis (figure S8-C). First, we counted the total number of recorded beads, N. Then, according to the normal distribution assumption, we drew the expected sigma count distribution along the z axis inside the range [-2.2 µm, 2.2 µm] with bin size equal to 10 nm. We fitted *σ*_*X*_, *σ*_*Y*_ histograms to the expected counts in z axis and obtained two calibration functions. During the localization stage of our pipeline, we used the mean estimation of z position of these two calibration functions.

Optimally, the PSF is the smallest when the cell is in focus and there is an equal number of beads above and below focus; however, the focus might change unintentionally over long acquisition periods leading to asymmetric empiric distribution. Therefore, during the calibration we divided the beads to two groups: horizontally stretched beads and vertically stretched beads. Then, we shifted the center of the Gaussian distribution to the real focal plane based on the ratio between horizontally stretched and vertically stretched PSFs.

#### 3D localization prediction in cells

We performed 3D localization analysis in the red and the green channels separately for each cell image. The localization process was composed of the following steps: 1) images were cropped to a constant size (25×25*pixels*^2^ or 7.5×7.5[μm^2^]); 2) images with bad SNR (SNR<15) were filtered out, where SNR was defined by the max intensity pixel value divided by the mean pixel value in each image; 3) each local maxima was fitted to a model, described in eq. 3, of an asymmetric Gaussian (modeling the PSF engineered locus) overlapping a rotated asymmetric Gaussian (modeling the cell background); 4) *x, y* position of the cells were extracted from the locus Gaussian fit parameters; 5) the predicted *σ*_*X*_, *σ*_*Y*_ parameters of the locus Gaussians were used to predict the z position.

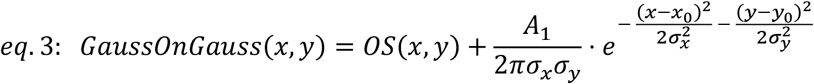

Where: *OS*(*x, y*) is the offset defined by the cell background (see eq. 4), *A*_1_ is the locus amplitude estimate *σ*_*X*_, *σ*_*Y*_ are the locus Gaussian standard deviation estimates, and *x*_0_, *y*_0_ are the locus location estimates.

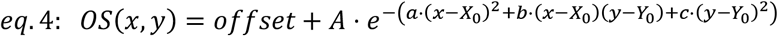

Where: *offset* is the mean background signal in the image, A is the cell amplitude estimate, *x*_0_, *y*_0_ are the cell location estimates, and a, b, c are the parameters of an asymmetric gaussian rotated by the angle *θ*:

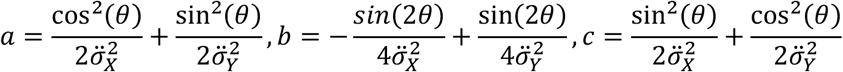

Where 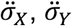 are the cell Gaussian standard deviation estimates.

If the *R*^2^ fit score was less than 0.9, we removed the locus from the analysis pipeline. Finally, we used the precalculated calibration functions for *σ*_*x*_ and *σ*_*y*_ and averaged their predictions to get the final estimate of the z position.

In cases where there were multiple loci in a single image, we validated that the number of loci detected was the same in both channels, otherwise the cells were filtered out from the analysis. Furthermore, we did not analyze loci that were located within distances of 5 pixels to avoid ambiguity due to extremely large PSFs.

The ImageStream device detects different cells in different positions due to flow rate variability over time and space. Namely, different cells are detected in different positions on the ImageStream sensor. To overcome biases in the predicted position due to the cell position relative to the sensor, we applied mean subtraction of the mean predicted x, y positions from each cell’s predicted x, y positions. Notably, this step substantially improved the performance of our pipeline, i.e. the bead distance histograms got closer to zero after applying this step.

Additionally, at each time point, all cell frames included in the analysis were manually reviewed to exclude any bead frames that were mistakenly incorporated.

#### 3D distance prediction

In this study, we reported the mean 3D distance values of the analyzed cell population at each time point. The 3D distances between the loci in the green channel and the red channel per cell was calculated by L_2_ distance (eq. 5):

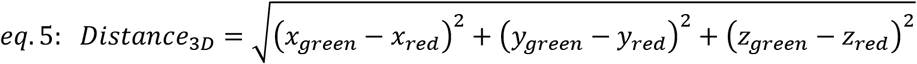

## Data availability

The data underlying this article will be shared upon reasonable request to the corresponding author.

## Code availability

The IFC analysis code is available at https://github.com/alonsaguy/DNA-double-strand-break-repair-quantification-and-tracking

## Supplementary data

Supplementary data are available online.

## Author contributions

Yael Shalev Ezra: Conceptualization, Methodology, Visualization, Data analysis, Writing— original draft. Alon Saguy: Data analysis. Lucien Weiss: Methodology. Onit Alalouf: Conceptualization, Supervision, Methodology, Writing. Yoav Shechtman: Conceptualization, Supervision, Methodology, Writing.

## Acknowledgements

We would like to thank Dr Elisa Dultz, Prof. Karsten Weis, Prof. Kerstin Bystricky, Prof. James R. Broach, Prof. James E. Haber, Prof. Nabieh Ayoub and Prof. Yoav Arava for providing us with plasmids, yeast strains and very valued advice and discussions.

We also would like to thank Maayan Duvshani, Yosuf Mansur, and Aviv Lutaty and the Life Science and Engineering (LS&E) Infrastructure Center for the technical assistance with the flow cytometry devices.

## Ethics declarations

Conflict of interests - The authors declare no conflict of interest.

