## SUPPLEMENTARY MATERIALS FILE for "High-throughput DNA repair monitoring in *Saccharomyces cerevisiae* reveals SSB- and DSB-induced chromatin reconfiguration"

### Table of figures

Figure S1: Inducible and reporting strain construction and mating-type switching quantification 7

### Table of tables

#### Supplementary materials

##### *S1. Validation of mating-type switching efficiencies*

To monitor the mating-type switching process in this study, we constructed *MATa-stk* strains containing inducible HO-endonuclease and a CFP or 6XmsCFP as a switching reporter. For switching induction, we grew the cells according to 'Mating type switching by galactose induction' in the methods section and obtained switching efficiencies of 5-20%.

These switching levels are considered to be low compared to the natural[1] and induced processes[2]. We checked if the *MATa-stk* mutation was the sole factor contributing to the low switching efficiencies by testing the following: **1.** Transformed plasmid 'Popout' from the genome (figure S2-A); **2.** Alternative DSB repair mechanisms relating to the population cell cycle phase shift due to non-fermentable sugar medium (figure S2-B-D); **3**. Re-switching during induction (figure S2-F); **4.** FROS labeling interference with the switching process (figure 4).

To check if the low switching efficiency was due to 'Popout' of the HO-CFP plasmids from the yeast genome, we grew raffinose starters to induction stage ($O.D_{600}$ ~0.3), plated diluted cultures (1000-fold) on YEPD plates, incubated the plates at 30°C for two nights and replica plated the colonies on SD plates without Methionine (the marker on the HO-CFP plasmid). We observed 100% growth on the selectable medium, which indicated that 'Popout' did not occur (figure S2-A).

We then evaluated the effect of the sugar used, raffinose, on the switching levels in our experiments. Naturally, mating type switching occurs mainly in the late G1 phase in the mother cell before DNA replication (S/G2 phase)[3]. We plotted the cell cycle stage distribution (G1 and S/G2) of the pre-induced populations by measuring DNA content in fixed nucleus-stained cells to determine the correlation between the medium, the cell cycle stage and repair by gene conversion (mating-type switching occurrence). The tested strains (BY4741, YS0075 (HO-CFP), YS0095 (HO-6xCFP), YS0099 (HO-CFP+ 2 FROSs)) had an ~1.5-fold more cells in G1 relative to S/G2 when grown in raffinose, and about twice as many cells in G1 than in S/G2 when grown in YEPD medium. Thus, the low switching efficiency in raffinose containing medium might be cell cycle related (figure S2-B). To investigate this point further we checked if higher switching efficiencies can be achieved by using YEPD starters rather than raffinose starters. We tried two strategies: 1. YEPD only - growing YEPD starters overnight, centrifuging the cells, and diluting them to $O.D_{600}$ ~0.3 in 20 ml raffinose, and immediately (30-60 min) starting the switching experiment; 2. YPD/raffinose - growing YEPD starters overnight, diluting the cells into the raffinose medium, growing the cultures overnight, and starting a switching experiment at $O.D_{600}$ ~0.3. In both cases, switching levels remained unchanged relative to the original raffinose starter case (figure S2-C-D), emphasizing minimal cell cycle influence. Importantly, we noticed that inadequate adaptation of the cultures to the raffinose medium significantly reduced the quality of FROS localizations, especially for the LacO-LacI-eGFP system (figure S2-E), making them unsuitable for imaging using PSF engineered image flow cytometry (IFC) or time-lapse microscopy.

The occurrence of re-switching led previous studies to use mutated strains, such as deletion of the HMRa[4] and mutating the HO-endonuclease recognition site in the HMLα (*HML-inc*)[5,6]. We determined the significance of re-switching in our strains by calculating switching efficiencies (by CFP reporter) with longer induction durations (2, 3 and 6, hours). Compared to a 1-hour induction switching experiment, induction of 2 and 3 hours resulted in increased switching efficiency of about 1.65% and 2.36%, respectively. We concluded that re-switching events were not significant and were not the reason for low switching levels. Interestingly after the induction of 6 hours, no switching was detected (figure S2-F). According to Haber *et al.*[6] and Xie *et al*.[7], in *MATa-stk* strains switching from 'α' to 'a' is more efficient than 'a' to 'α', which might suggest that re-switching prevailed[8]. However, we still expected to have some 'α' in the population if switching kept occurring. Another explanation is continuous switching induction that results in HMLα and HMRa cleavage[3], reducing the switching level. Lastly, we showed in the main text that FROS integration into the yeast genome reduced switching efficiencies by 2-3-fold.

In summary, the low switching efficiencies in our experiments were mainly due to inefficient DSB by the HO-endonuclease due to the *MATa-stk* mutation in the BY4741 derivatives[8,9] and genome perturbation by the integrated FROS.

#### ***S2. Supplementary methods (used in section S1)***

##### ***I. Viability test, mating type switching validation and 'Popout' assay***

Determining culture mating type by plate assay requires crossing the culture with tester strains of known type with auxothrophic marker. Tester strains 17-14 (MATa, *-his*) and 17-17 (MATα, *-his*)[7] were grown to stationary phase, 200 µl from each culture were spread on YEPD plates and were incubated at 30°C overnight for confluent layer.

Cultures post mating-type switching experiments (induced and uninduced) were diluted by 1000, and 50 µl of each were spread on YEPD plates and incubated at 30°C for 1-2 days. Colonies on each plate were counted for viability, $\%\ viability = 100 * \frac{\mathrm{N_{cells\_induced}}}{N_{\mathrm{cells\_uninduced}}}$, crossed with the tester strains by replica plating on a fresh YEPD plate and incubated at 30°C for 1-2 days. The YEPD plates were then replicated to the appropriate SD plates and incubated at 30°C for additional 1-2 days to select for dipolds colonies (indicating switching).

For the ‘Popout’ assay, YEPD plates with post-switching diluted cultures were incubated as above, replicated to the SD(-Met) plates and incubated at 30$^{o}$C for additional 1-2 days to select for colonies that did not discard the HO-CFP plasmid.

##### *II. Cell cycle assay by flow cytometry*

To assess the cell cycle stages prior to mating type switching induction, cultures were grown according to ‘Mating type switching by galactose induction’ in the methods section. Cultures reaching $O.D_{600}$ ~0.3 were fixed and their DNA stained as described in Harari *et al*., 2018[10] with a few modifications: 400 µl aliquotes were centrifuged at 8,000-13,000 rcf for 1.5 minutes, and the pellets washed with 200 µl TE solution (10mM TrisHCl pH 7.5, 1mM EDTA) and fixed overnight at 4ºC with 600 µl of 70% ethanol in TE buffer. Cells were harvested and washed twice with 400 µl cold TE buffer, re-suspended in 200 µl of TE-RNaseA solution (0.25mg/ml RNaseA), and incubated for 2 hours at 37 $^{o}$C. Cells were harvested and washed using 400 µl cold TE buffer, re-suspended in 200 µl of TE-proteinase-K solution (0.25 mg/ml proteinase-K), and incubated for 1 hour at 50$^{o}$C. Cells were then harvested and washed using 400 µl cold TE buffer, re-suspended in 200 µl of TE-PI solution (20 mg/ml Propidium-iodide) and incubated overnight at 4ºC and in the dark. The fixed and stained cells were loaded into a 96-well plate, sonicated 3 times for 2 seconds at 50J power, and verified to be singlets under the microscope (no clusters). The samples were then analyzed using the Cytek Aurora full spectral FC device and the SpectroFlo software.

#### *S3. Supplementary results*

##### *I. Tracking DNA dynamics continuously in mating-type switching induced culture by Imaging flow cytometry*

To observe the HMLα-MAT loci distances with higher temporal resolution during switching we acquired images continuously for 3 hours (rather than every 30 minutes) using the IFC (figure S6). We then analyzed 3-minute windows and observed similar behavior to the 30-minute intervals, but at higher temporal resolution: loci proximity was detected sooner, after 10 minutes from induction (in agreement with Hicks *et al.*[4], which reported DSB 10 minutes after induction in 50% of the population), with an average distance shortening of ~100-150 nm between the loci (relative to 250 nm at t=0 minutes and 300 nm in the uninduced culture) remained up to 2 hours (in agreement with Bressen *et al.*[11]; figure S6).

#### Supplementary figures

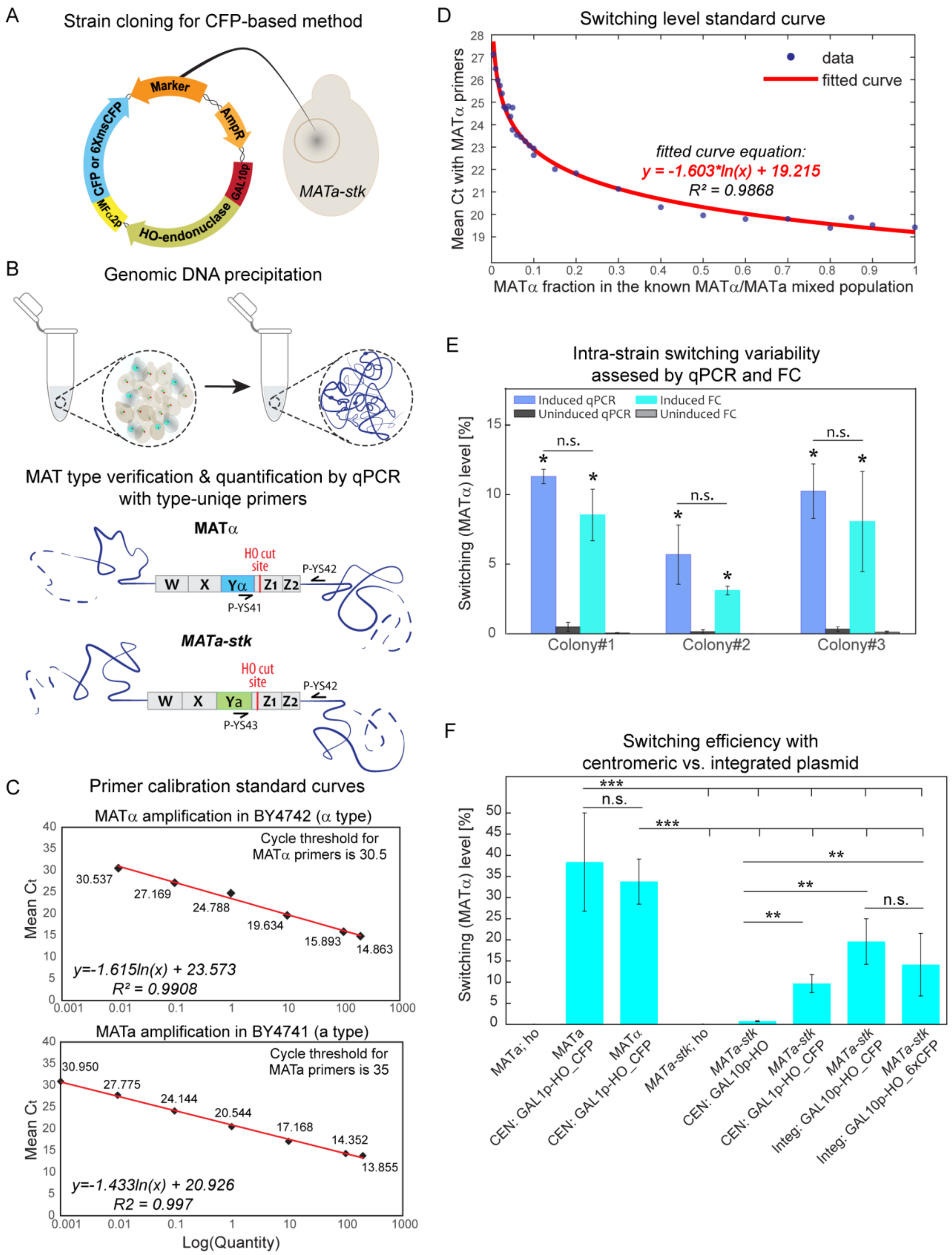

**Figure S1: Inducible and reporting strain construction and mating-type switching quantification** (A) Construction of a HO-CFP or HO-6XCFP mating-type switching *MATa-stk* strains, stably expressing the HO-endonuclease under the inducible GAL10 promoter and CFP (YS0075) or 6XmsCFP (YS0095), respectively, under the MFα2 promoter. (B) Schematic illustration showing the qPCR process starting from genomic DNA precipitation to the qPCR of the MATα or MATa loci using either the PYS-41 or PYS-43 primers, respectively, together with PYS-42 primer. Damaged DNA region cannot be amplified. (C) calibration curves for the PYS-41 and PYS-42 primers in BY4742 (MATα) and BY4741 (*MATa-stk*), determining the cycle threshold (Ct) at 30.5; calibration curves for the PYS-43 and PYS-42 primers in BY4741 (*MATa-stk*), determining the Ct at 35. (D) Population switching level was determined based on qPCR standard curve with MATα primers (PYS-41 PYS-42) and known BY4741 (*MATa-stk*)/ BY4742 (MATα) mixtures in the range of 0.1-100% BY4742 (MATα) (detection limit for this assay was 0.1%). (E) Switching levels in YS0095 (HO-6XCFP) 6 hours after induction were measured both by qPCR and flow cytometry (switching level differences were significant between induced and uninduced cultures (p-value =0.0001), between colonies (p-value=0.0017), and non-significant (n.s.) between the two methods (p-value =0.2753)). (F) Switching efficiencies of the HO-CFP and HO-6XCFP systems were compared to two centromeric plasmids (pJH132[2] and B2609[12]) in BY4741 (*MATa-stk*), BY4742 (MATα; WT) and W303-1A (MATa; WT) strains, using both the CFP reporter (where applicable) and qPCR for switching quantification. Switching levels in WT strains, either MATa or MATα, were up to 40-fold higher compared to the *MATa-stk* strains. However, within the *MATa-stk* strains, HO-CFP and HO-6XCFP plasmids resulted in up to 10-fold higher levels of switching, relative to the centromeric plasmids. All experiments were done with at least 2 biological replicates.

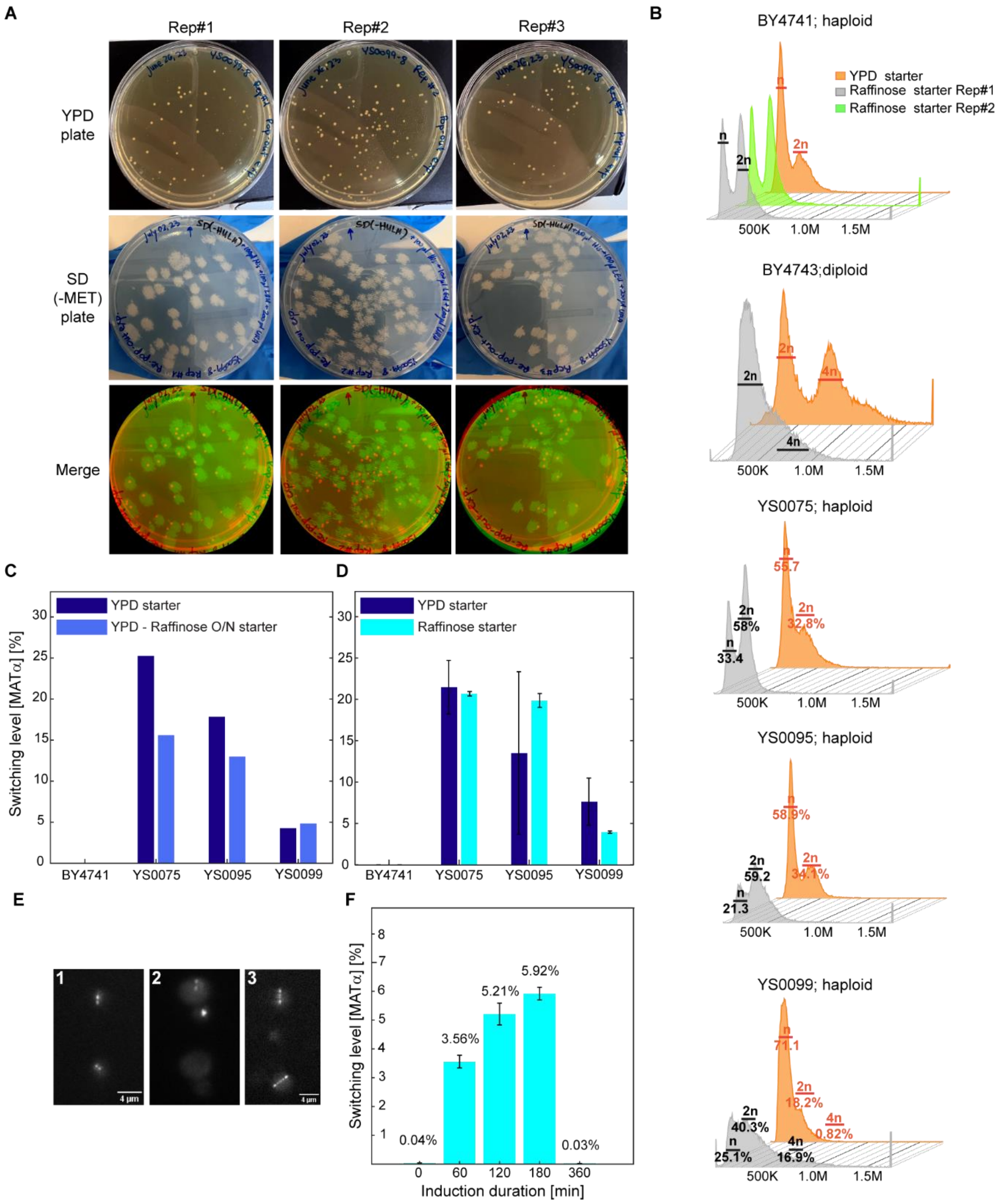

**Figure S2: Mating-type switching validation assay.** (A) replica plate assay for 'Popout' analysis (see supplementary methods S2-I). (B) Effect of raffinose on the cell cycle stage distribution (G1 and S/G2). DNA content increased ~1.5-fold in S/G2 when grown on raffinose, compared to YPD. (C-D) The effect of starter growth media on induced switching levels: (C) YEPD starter versus YEPD/raffinose starter, and (D) YEPD/raffinose starter versus raffinose starter. (E) Cells grown in (1) YEPD have localizations with higher SNR compared to, (2) cells that are grown overnight in raffinose with no adaptation to the medium (especially for the LacO-LacI-eGFP). (3) Adaptation to the raffinose medium (by 2-7 days incubation, depending on the strain) restores the localization SNR. Cells were imaged using a PSF engineered optical setup[13]; the mCherry and eGFP localizations were spectrally separated to the upper and lower parts of the same image, respectively. (F) The Effect of induction duration on switching levels in the dually tagged strain (YS0099).

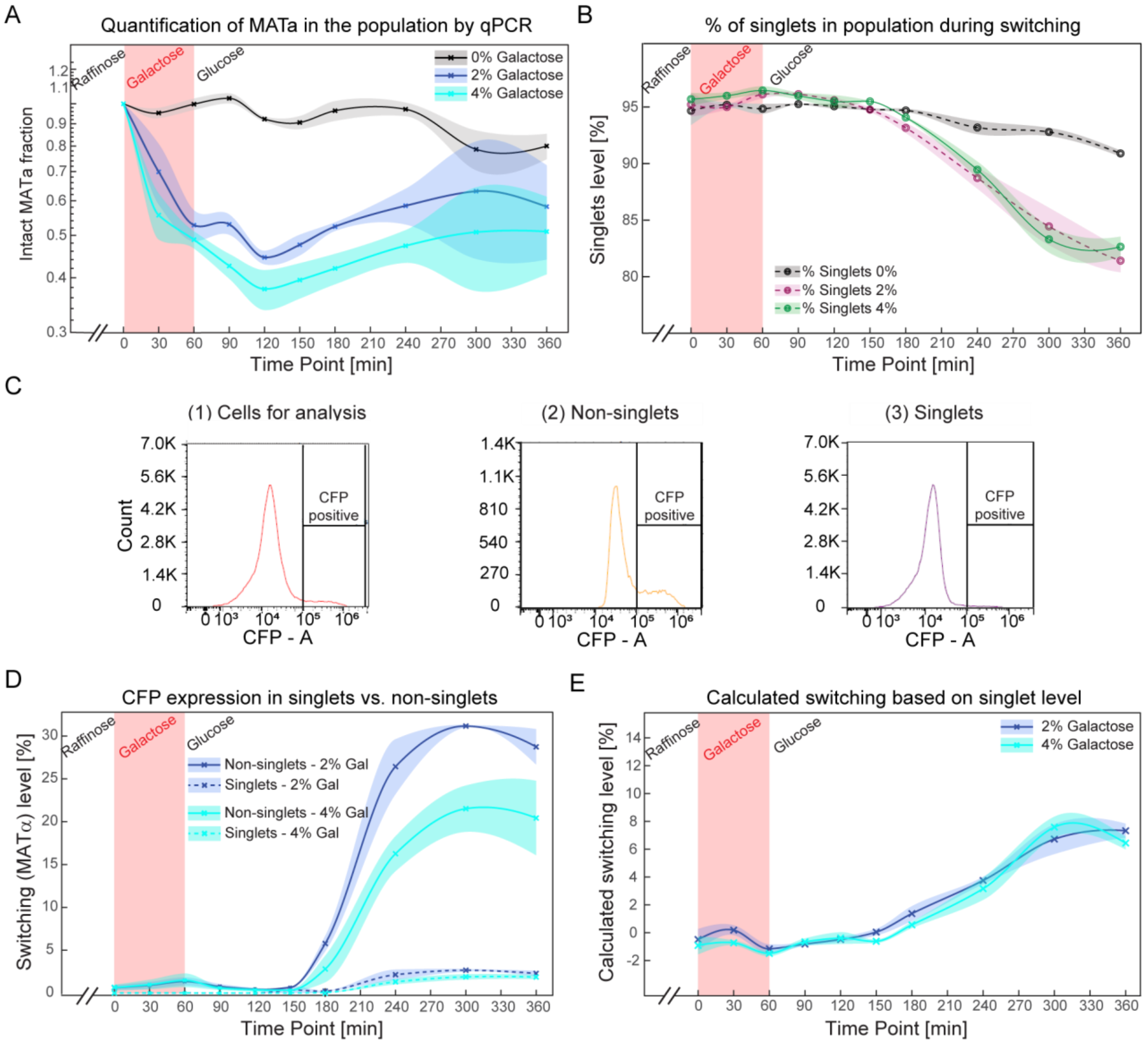

**Figure S3**: **Real-time quantification of DNA damage and mating-type switching.** After switching induction (A) *MATa-stk* locus amplification by qPCR is reduced and negatively correlates with the amount of DNA damage (DNA polymerase fails to amplify cleaved DNA); and (B) Single cells (singlets) number is reduced over time as seen by flow cytometry (see figure S7). (C, D) Switched cells appear mainly as non-singlets in flow cytometry due to mating events; (C) sample gating for switching quantification by CFP from single time-point in (1) cells for analysis (2) non-singlets and (3) singlets. (D) This was validated by measuring and comparing the CFP levels (switched cells) in the singlets and the non-singlets in each time-point. (E) Calculated switching levels from singlet levels (obtained using eq. 1). Singlet levels are negatively correlated with switching levels and can be used as a label-free method to calculate mating type switching in *MATa-stk* strain.

Table S1: Features of mating-type switching quantification methods developed in this study

| **Feature/Method** | **qPCR** | **CFP-based method** | **Label-free (flow cytometry)** |
|---|---|---|---|
| **System preparation** | 1. Gal-HO plasmid design and cloning<br>2. Transformation<br>3. Design of mating-type specific primers | 1. Gal-HO and CFP mating type specific plasmid design and cloning<br>2. Transformation | 1. Gal-HO plasmid design and cloning<br>2. Transformation |
| **Equipment** | qPCR system | Flow cytometry / Fluorescence microscopy | Flow cytometry |
| **System calibration** | Yes | No | No |
| **Culture type** | Suspension | Suspension or immobilized cells | Suspension |
| **Sample preparation/ processing time** | Genomic DNA extraction and qPCR reaction preparation (~2 days*) | Does not require special sample preparation | Does not require special sample preparation |
| **On-line tracking of switching levels** | No | Yes | Yes |
| **a-to-α switching-detection lag-time** | <30 minutes Depending on the time interval between the measurements | 150 minutes (Suspension) Depending on CFP expression and maturation time[14] | 120-150 minutes Depending on MATα protein expression |
| **Sensitivity** | 0.1% in the population | 1 cell | 1 cell |

* Based on 3 biological replicates of 10 time points.

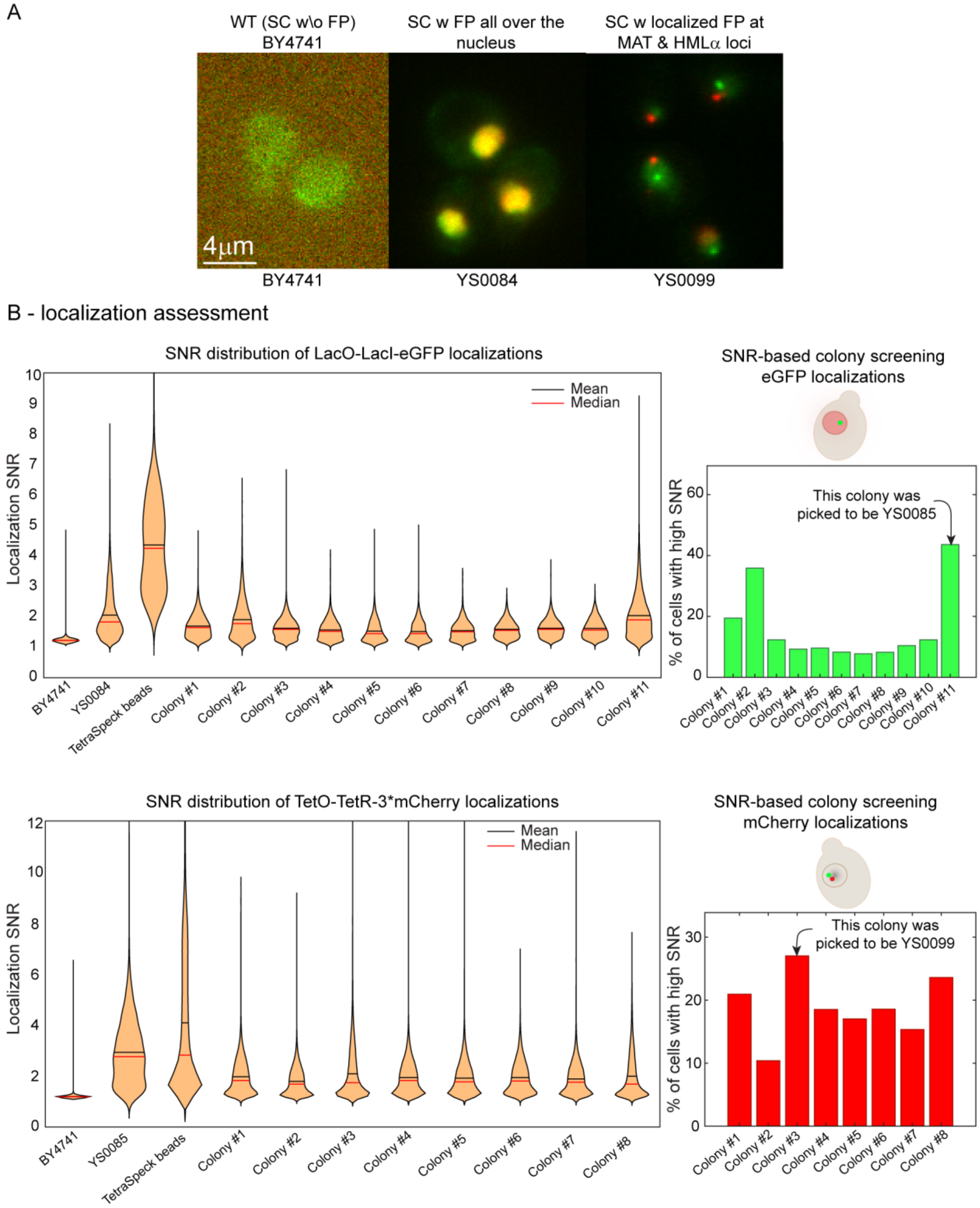

**Figure S4: Construction of *MATa-stk* and HMLα tagged strain.** (A) Representative widefield microscopy images of BY4741 strain (without fluorescent proteins); YS0084 strain expressing LacI-eGFP and TetR-3XmCherry in the nucleus; and YS0099 strain with two FROSs, 256 LacO repeats integrated proximal to the *MATa-stk* locus and 112 TetO repeats integrated proximal to the HMLα locus. (B) The dually tagged strain construction was sequential; initially transforming YS0084 with the 256 LacO repeats yielding the YS0085 strain, followed by transforming YS0085 with the 112 TetO repeats yielding the YS0099 strain. In each step, at least 8 colonies were scanned using the IFC device and selected based on the SNR value. WT – wild type; SC – *S. cerevisiae*; W – with; W/O – without; FP – fluorescent protein; SNR – signal to noise ratio..

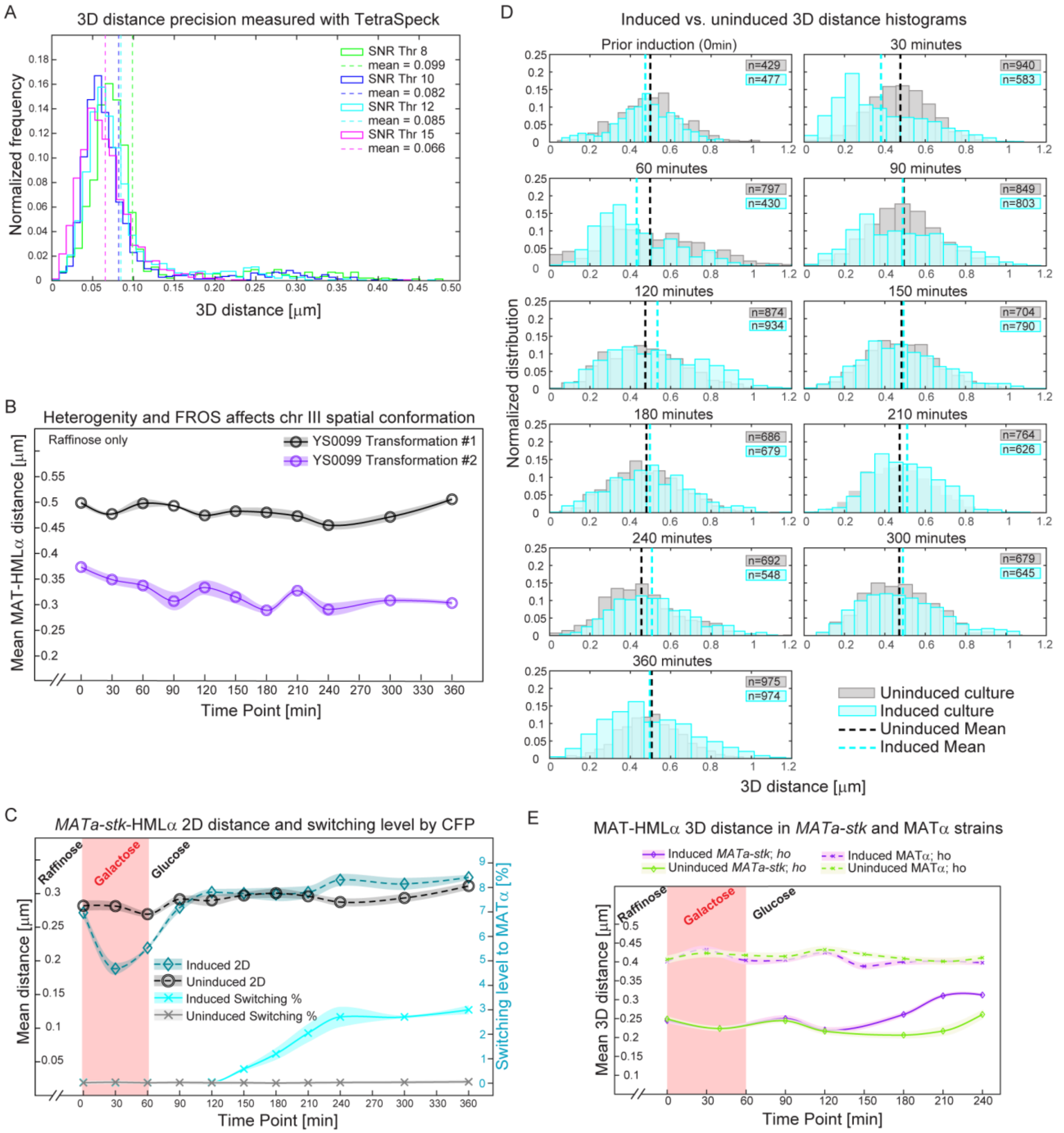

**Figure S5: Visualization of *MATa-stk* and HMLα loci distance dynamics during switching with IFC.** (A) Multicolor fluorescent TetraSpeck beads (0.2 μm) were used to assess the IFC method precision with different SNR thresholds (8, 10, 12 and 15). We chose SNR threshold of

15, that gave 60-80 nm precision and at least 100 analyzable cells at each time point for each replicate. (B) 3D distance heterogeneity between the MAT and HMLα loci in a strain that was obtained by different transformations with the same plasmids, illustrating transformation-dependent chromatin configuration. (C) Mean 2D distance between *MATa-stk* and HMLα and switching levels by CFP detection of an induced and uninduced cultures (3 replicates were gathered and analyzed via bootstrapping: mean (line) ± s.d. (shaded region representing the estimated error) of eight independent groups at each time point, overall >429 cells per time point). Distance shortening and switching were detected at 30 and 150 minutes, respectively, post switching induction (p-value << 0.05). (D) Histograms of real time 3D distance measurements between the *MATa-stk* and HMLα loci, for each time point the numbers of analyzed cells is written for both the induced and uninduced cultures. (E) The distance between the *MATa-stk* and HMLα loci is not influenced by the galactose addition, and is mating-type dependent (using unswitchable *MATa-stk* and MATα strains). Typically, the HMLα in MATa cells is closer to MAT locus than in the MATα cells[15].

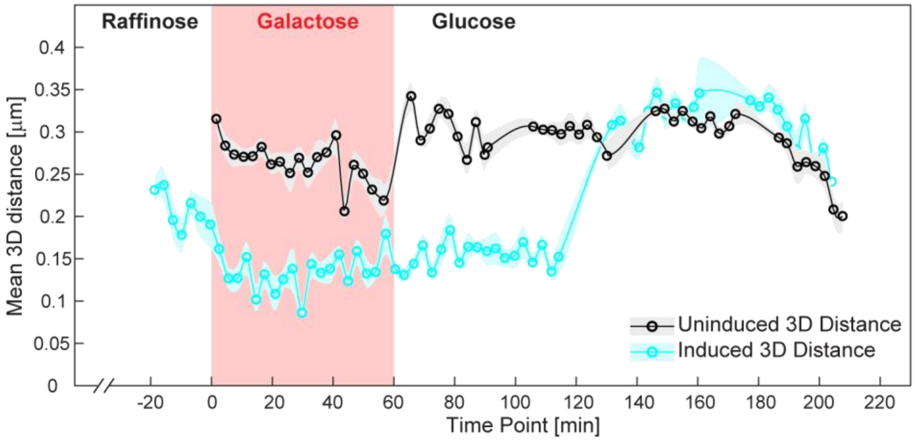

**Figure S6: Continuous *MATa-stk*-HMLα 3D distance tracking during mating-type switching.** Real-time distances between the *MATa-stk* and HMLα loci during switching was assessed continuously for 3 hours, both in an induced and uninduced cultures. The 3D distances were divided and analyzed as 3-minute windows by bootstrapping: mean (line) ± s.d. (shaded region representing the estimated error) of eight independent groups at each time point with hundreds of cells in each frame.

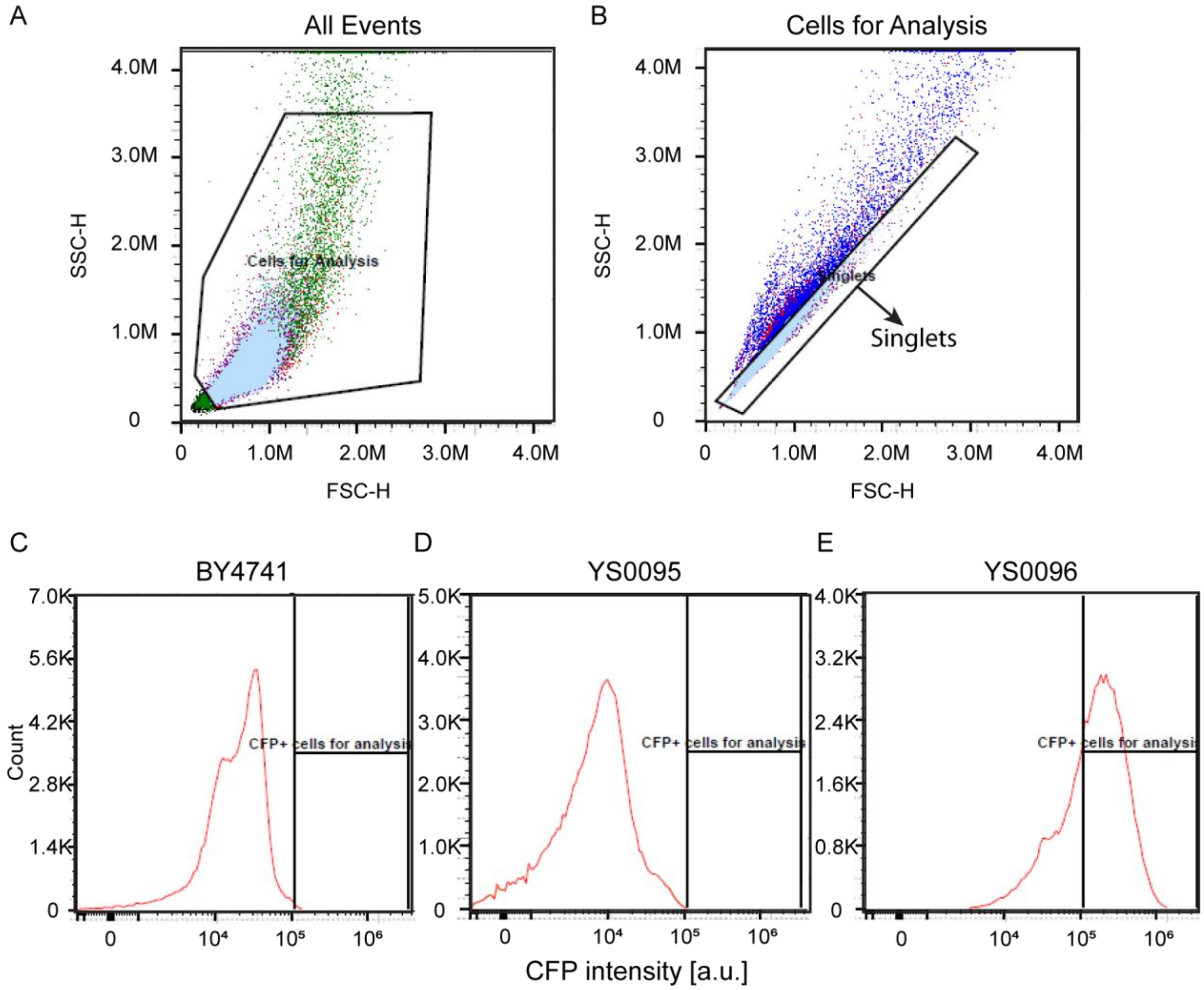

**Figure S7: Flow cytometry gating for the detection of CFP expressing cells.** (A) Cells for analysis were gated based on side and forward scattering (SSC-H x FSC-H) parameters, discarding debris and big aggregates, followed by (B) singlets that were gated based on their rounded shape. (C-E) Switched cells within the culture were gated based on CFP intensity, using BY4741 (unstained uninducible strain) and YS0095 (unstained strain prior to induction) as background strains, and YS0096 as a positive control expressing CFP constitutively.

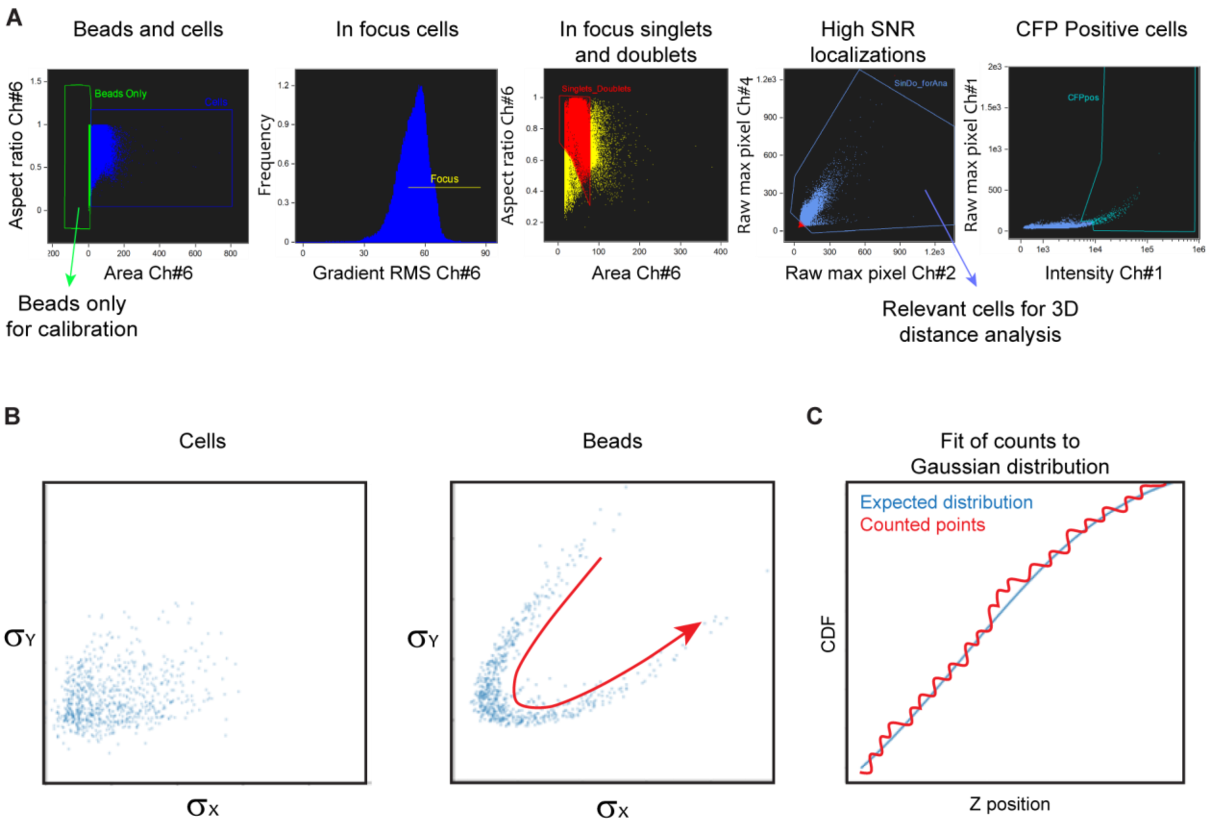

**Figure S8: IFC data acquisition and calibration.** (A) Beads and cells were separately gated based on their area versus aspect ratio in the brightfield channel. Gradient RMS further filtered the cells for in-focus, gated for singlets or doublets, and those with high SNR in both loci (eGPF and mCherry) were analyzed. Population switching levels were calculated according to the CFP signal at each time point. (B) For PSF calibration we used TetraSpeck beads, rather than cells due to background noise and low SNR, to calculate the Gaussian standard deviation estimates ($\sigma_X$, $\sigma_Y$) and (C) to fit the position to the expected Gaussian distribution (see methods for more details). Data files were exported as separate cif and txt files for calibration and analysis. CDF – cumulative distribution function.

#### **Supplementary tables**

Table S2: plasmid used in this study.

| Plasmid name | Property | Resistance, Copy Number | Yeast homologous recombination site | Marker for integration | Restriction enzyme for linearization | Reference |
|---|---|---|---|---|---|---|
| pUC19 | Backbone | Amp, H | NA | NA | NA | |
| pKW3034 | HISp: LacI-eGFP; URAp: TetR-3*mCherry; | Amp, H | Chromosome XV (his3Δ1) | HIS3 | NheI | Dultz *et al*. 2018[16] |
| pKW1689 | 256 LacO repeats | Amp, H | Chromosome II (YBR022 - POA CDS) | LEU2 | BglII | Dultz *et al*. 2018[16] |
| pKW2837 | 112 TetO repeats | Amp, H | Chromosome II (YBR022 - POA CDS) | URA3 | BlpI | Dultz *et al*. 2018[16] |
| pIL01 | HISp: LacI-CFP; URAp:λI-YFP; ADH1t; | Amp, H | NA | NA | NA | Lassadi *et al*. 2015[17] |
| pCyPet-His | pCyPet-His | CM, H | NA | NA | NA | Nguyen *et al*. 2005[18] |
| BS-Met15 | BS-Met15 | Amp, H | NA | NA | NA | Li D *et al*. 2015[19] |
| pJH132 | GAL10p-HO, URA3, CEN | Amp, H | NA | NA | NA | Wu & Haber, 1995[2] |
| B2608 | LEU2 CEN4 ARS1 PGAL1-HO PMFa1-CFP | Amp, H | NA | NA | NA | Houston & Broach, 2006[12] |
| B2609 | LEU2 CEN4 ARS1 PGAL1-HO PMFα1-CFP | Amp, H | NA | NA | NA | Houston & Broach, 2006[12] |
| pYS0019 | 112 TetO repeats | Amp, H | Chromosome III (15088-15621bp; HMLα proximal) | URA3 | BmgBI | This study |
| pYS0020 | 256 LacO repeats | Amp, H | Chromosome III (15088-15621bp; HMLα proximal) | LEU2 | SpeI | This study |
| pYS0021 | 256 LacO repeats | Amp, H | Chromosome III (197,202-197,392bp; MAT proximal) | LEU2 | PmlI | This study |
| pYS0080 | GAL1p-HO; MFα2p-CyPet | Amp, H | Chromosome XII (730,361-730,807bp) | MET15 | AflII | This study |
| pYS0088 | GAL1p-HO; MFα2p-CFP | Amp, H | Chromosome XII (730,361-730,807bp) | MET15 | AflII | This study |
| pYS0099 | GAL10p-HO; MFα2p-CFP | Amp, H | Chromosome XII (730,361-730,807bp) | MET15 | AflII | This study |
| pYS0107 | GAL10p-HO; MFα2p-6XmsCFP | Amp, H | Chromosome XII (730,361-730,807bp) | MET15 | AflII | This study |

Table S3: primers used in this study.

| **Primer #** | **Sequence (5' to 3')** | **Target plasmid** | **Comments** |
|---|---|---|---|
| P-YS1 | GTCGACTCTAGAGGATCCCCGTTGATCTGCCTTTTATAGCTAAG | pYS0080 | |
| P-YS2 | CCGCCGAATAATTCTTCACCTTTAGACATTTTCTCCAATATGTGAATTTACTGG | pYS0080 | |
| P-YS3 | CCAGTAAATTCACATATTGGAGAAAATGTCTAAAGGTGAAGAATTATTCGGC | pYS0080 | |
| P-YS4 | CTGGCGAAGAAGTCCAAAGCTTTTGTACAATTCATCCATACCATGGG | pYS0080 | |
| P-YS5 | CCCATGGTATGGATGAATTGTACAAAAGCTTTGGACTTCTTCGCCAG | pYS0080 | |
| P-YS6 | GTGAATTCGAGCTCGGTACCTAGAGGATCCGTGTGGAAGAACG | pYS0080 | |
| P-YS7 | CACACCGCATATGGTGCACTCTCAGTACAATCTGTCATGGTTTTTGGCCAGCG | pYS0080 | |
| P-YS8 | CGGGGCTGGCTTAACTATGCGGCATCAGAGGGTTCGAATCCCTTAGCTCTC | pYS0080 | |
| P-YS9 | CTGTTTTCGCTGGCCAAAAACCATGACAGATTGTACTGAGAGTGCACC | pYS0080 | |
| P-YS10 | GAGAGCTAAGGGATTCGAACCCTCTGATGCCGCATAGTTAAGC | pYS0080 | |
| P-YS11 | CACTCATTAGGCACCCCAGGCTTTACACCAGTCCGCTCATTTTAGCTG | pYS0080 | |
| P-YS12 | CCACACAACATACGAGCCGGAAGCATAAACCAGTTACAACAGCGGTGAG | pYS0080 | |
| P-YS13 | CTACCATCACCTGCATCAAATTCCAGTAAATTCACATATTGGAGAAAATGAGTAAAGGAGAAGAACTTTTCACTG | pYS0088 | |
| P-YS14 | GCTTTAGGCAACCTTTCTCTTCTTCTTTGGTGGAGTACAGGATCCCAGTTTGTATAGTTCATCCATGCCATGTG | pYS0088 | |
| P-YS15 | CAGGAAACAGCTATGACCATGATTACGCCAAGCTTCCTATAAAAATAGGCGTATCACGAGGC | pYS0099 | |
| P-YS16 | CGGGGATCCTCTAGAGTCGACGAAGAGAGGTTCCAAGTCCAAGATTG | pYS0099 | |
| P-YS17 | CCAGCGGGTAAAG**CGGCCG**AGCTTTGGACTTCTTCGCCAG | pYS0107 | EagI restriction site on primer (bold red) |
| P-YS18 | GTGCTATCCAT**GCTAGC**TGGTGGGGCAACCTTTCTCTTCTTCTTTGGCATTTTCTCCAATATGTGAATTTACTGG | pYS0107 | NheI restriction site on primer (bold red) |
| P-YS19 | TTGCATACAAATGAC**GAGCTC**CGAGGGGCTTTACTGTAATATATG | pYS0019 | SacI restriction site on primer (bold red) |
| P-YS20 | CTCAATAGACATACAGTGCGGGCATCAATTTGCCCTTGATAATTTTC | pYS0019 | Restriction site for NgoMIV is not on the primer (on HMLα proximal) |
| P-YS21 | GGTCATGGCATTAATCCTA**GGTACC**TTTAAATGTTGAGGTAAATAGC | pYS0020 | Acc65I restriction site on primer (bold red) |

Table S3 continue: primers used in this study.

| Primer # | Sequence | Target plasmid | Comments |
|---|---|---|---|
| P-YS22 | CGAGTTCATGATAGGACATTGGAAGTAATCTGCAGATGCGG | pYS0020 | Restriction site for XhoI is not on the primer (on HMR proximal) |
| P-YS23 | GACCTATGCATG**CTCGAG**CCGCCTTTACCGTAGTTTTG | pYS0021 | XhoI restriction site on primer (bold red) |
| P-YS24 | CATCGATCAGCAAT**GGTACC**GTATCTTTAGGCTACCGC | pYS0021 | Acc65I restriction site on primer (bold red) |
| P-YS25 | GGGATGCTCAAAGATTGATGTTCAATGGTTTATTCCCTC | pYS0099 & pYS0107 | Upstream integration verification by colony PCR |
| P-YS26 | GTTTGGTAGGGACCCTGACTGAC | pYS0099 & pYS0107 | Upstream integration verification by colony PCR |
| P-YS27 | GCCTCACTGATTAAGCATTGGTAACTGTCAGACCAAG | pYS0099 & pYS0107 | Downstream integration verification by colony PCR |
| P-YS28 | GGTGAATGTTCTTTCGTTTGGATGAGGACCCAC | pYS0099 & pYS0107 | Downstream integration verification by colony PCR |
| P-YS29 | CTACTCGGGTTCAGCAACGCTG | pKW3034 | Upstream integration verification by colony PCR |
| P-YS30 | CTGTGCGGTATTTCACACCGC | pKW3034 | Upstream integration verification by colony PCR |
| P-YS31 | GTCTGTAAGCGGATGCCGGG | pKW3034 | Downstream integration verification by colony PCR |
| P-YS32 | CACTTGTTCGCTCAGTTCAGCC | pKW3034 | Downstream integration verification by colony PCR |
| P-YS33 | GCGGGAGCTTTCTTTCGATCTC | pYS0019 | Upstream integration verification by colony PCR |
| P-YS34 | CCGTCGAACCATCACCCTAATC | pYS0019 | Upstream integration verification by colony PCR |
| P-YS35 | GGATCCATACTAATCGGATCCCCG | pYS0019 | Downstream integration verification by colony PCR |
| P-YS36 | GGCTCAAAATTAGCTGCGGATGAC | pYS0019 | Downstream integration verification by colony PCR |
| P-YS37 | CTGAGAGTCAGCGTCATCGT | pYS0021 | Upstream integration verification by colony PCR |
| P-YS38 | CAACTGTTGGGAAGGGCGAT | pYS0021 | Upstream integration verification by colony PCR |
| P-YS39 | ATTTGTGGAATTCTCGAGCCG | pYS0021 | Downstream integration verification by colony PCR |
| P-YS40 | CGACAACAAAGACAGGGCGT | pYS0021 | Downstream integration verification by colony PCR |
| P-YS41 | GCAGCACGGAATATGGGACT | NA | |
| P-YS42 | ATGTGAACCGCATGGGCAGT | NA | |
| P-YS43 | GGCATTACTCCACTTCAAGT | NA | |
| P-YS44 | GATTCTGAGGTTGCTGCTTTG | NA | |
| P-YS45 | ACCGACGATAGATGGGAAGA | NA | |

Table S4: strains used and constructed in this study.

| Strain name | Genotype | Reference |
|---|---|---|
| BY4741 | MATa his3Δ1 leu2Δ0 met15Δ0 ura3Δ0 | Brachmann *et al*. 1998[20] |
| BY4742 | MATα his3Δ1 leu2dΔ0 lys2Δ0 ura3Δ0 | Brachmann *et al*. 1998[20] |
| W303-1A | MATa ade2-1 ura3-1 his3-11 trp1-1 leu2-3 leu2-112 can1-100 | *Saccharomyces cerevisiae* Meyen ex E.C. Hansen (ATCC 208352) |
| 17-14 | MATa -his | Ze-Xiong Xie *et al*., 2018[7] |
| 17-17 | MATα -his | Ze-Xiong Xie *et al*. 2018[7] |
| YS0057 | MATα his3Δ1::pKW3034 leu2Δ0 lys2Δ0 ura3Δ0 HMLα (15088-15621bp)::pYS0019 MAT(197,202-197,392bp)::pYS0021 | This study |
| YS0075 | MATa his3Δ1 leu2Δ0 met15Δ0 ura3Δ0 Chr XII (730361-730807bp position S288C)::pYS0099 | This study |
| YS0084 | MATa his3Δ1::pKW3034 leu2Δ0 met15Δ0 ura3Δ0 Chr XII (730361-730807bp)::pYS0099 | This study |
| YS0085 | MATa his3Δ1::pKW3034 leu2Δ0 met15Δ0 ura3Δ0 Chr XII (730361-730807bp)::pYS0099 MAT (197,202-197,392bp)::pYS0021 | This study |
| YS0095 | MATa his3Δ1 leu2Δ0 met15Δ0 ura3Δ0 Chr XII (730361-730807bp position S288C)::pYS0107 | This study |
| YS0096 | MATa his3Δ1 leu2Δ0 met15Δ0 ura3Δ0 Chr XII (730361-730807bp position S288C)::pYS0107 | This study |
| YS0097 | MATa/MATa his3Δ1 leu2Δ0 met15Δ0 ura3Δ0 Chr XII (730361-730807bp position S288C)::pYS0107 | This study |
| YS0087 / YS0099 | MATa his3Δ1::pKW3034 leu2Δ0 met15Δ0 ura3Δ0 Chr XII (730361-730807bp)::pYS0099 MAT (197,202-197,392bp)::pYS0021 HMLα (15088-15621bp)::pYS0019 | This study |
| YS0100 | MATa his3Δ1::pKW3034 leu2Δ0 met15Δ0 ura3Δ0 Chr XII (730361-730807bp position S288C)::pYS0099 HMLα (15088-15621bp)::pYS0019 | This study |
| YS0104 | MATa his3Δ1::pKW3034 leu2Δ0 met15Δ0 ura3Δ0 HMLα (15088-15621bp)::pYS0019 MAT(197,202-197,392bp)::pYS0021 | This study |
